# Dogs gone wild: habitat use and ecological impacts of feral dogs in sub-Antarctic Chile

**DOI:** 10.1101/2025.05.08.652634

**Authors:** Hana Tebelmann, Alicia Sophie Pages, Simon Käfer, Aintzane Cariñanos Ruiz, Nicolàs Soto Volkart, Udo Gansloßer

## Abstract

Feral dogs *(Canis familiaris)* are an emerging threat to biodiversity on Navarino Island, Chile, where they have become apex predators in the absence of natural carnivores. This study evaluated the spatial distribution of feral dogs and their impacts on native species, including guanacos *(Lama guanicoe)*, upland geese *(Chloephaga picta)*, and flightless steamer ducks *(Tachyeres pteneres)*. Presence-only data collected during two field expeditions were analysed using species distribution models (MaxEnt) to predict habitat suitability for feral dogs and guanacos. Habitat connectivity analyses identified at least two potentially isolated feral dog populations. Using generalised linear and non-linear models, we assessed the ecological impacts of feral dogs, finding significant habitat overlap with guanacos, particularly in central areas of the island. This overlap corresponded to a reduced likelihood of guanaco occurrence, suggesting behavioural adaptations to disturbance and predation pressure. Upland geese exhibited a negative association with both actual and predicted feral dog presence, while flightless steamer ducks appeared unaffected. Furthermore, NDVI changes observed on Navarino Island compared to Torres del Paine, where no invasive species are present, are likely linked to the spread of invasive species and the decline of guanacos in the Magellanic forest, highlighting the cascading ecological consequences of feral dog invasion. Our findings emphasise the urgent need for feral dog management to protect vulnerable species and maintain ecosystem health, particularly in fragile environments where invasive predators can have disproportionate impacts.

**Graphical Abstract:** 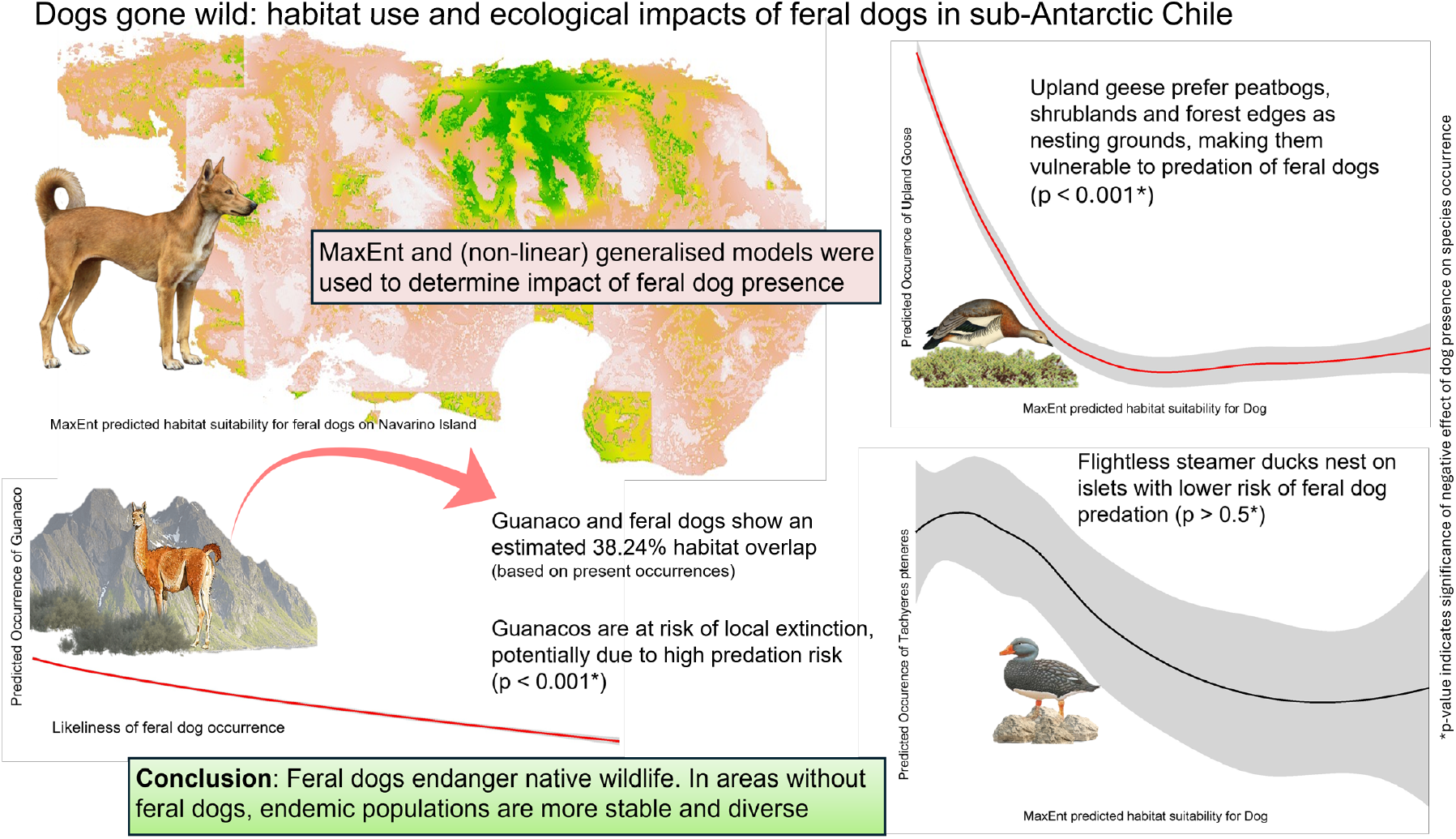

**Highlights:**

- Invasive species presence leads to environmental changes and potential tipping points
- Feral dogs pose major harm as an invasive species
- Endemic wildlife adapts their behaviour such as habitat use to avoid dog predation
- Species diversity and population stability are higher in areas with no feral dogs
- Dogs in general should be removed from the wild

## 1. Introduction

Biological invasions are increasingly prevalent and have become a significant driver of biodiversity loss [1, 2]. Invasive species are now found across virtually all ecosystems, including remote regions [3]. Among these, the domestic dog (*Canis familiaris*), with an estimated population of up to 987 million, stands out as the most abundant carnivore globally [4]. As an invasive species, dogs have a substantial impact on wildlife through predation and competition [5, 6, 7, 8].

Dogs affect wildlife in several ways, including altering spatial use [9], shifting temporal activity patterns [10], and increasing vigilance behaviours [11]. These behavioural changes likely impose considerable energetic costs on native prey species [12]. Mammalian predators are likely to create a landscape of fear in näıve endemic prey animals, causing potential maladaptive avoidance strategies and alternance in habitat use [13]. These behavioural adaptations can even cause trophic cascades, where changes in prey behaviour affect vegetation and other ecological processes [14]. This impact is particularly severe on islands, where the absence of natural predators and the lack of adaptive defenses in endemic species exacerbate their effects [15, 16]. Island species often evolve without facing competition or predation pressures, making them especially vulnerable [17].

On Navarino Island, located within the Cape Horn Biosphere Reserve (CHBR), recent studies have documented the presence of a feral dog population [18, 19]. This is particularly concerning as Navarino Island lacks native mammal predators, positioning feral dogs as apex predators. The native terrestrial vertebrate fauna consists of five mammal species, one freshwater fish species, and over 150 bird species [20, 21]. Notably, the number of invasive mammal species, totaling 12, surpasses that of native mammals, with dogs, cats, and American mink being among the terrestrial predators, and beavers (*Castor canadensis*) having additional severe impacts on the island’s ecosystem. Potential impacts of invasive predators include predation, such as feral dog predation on upland geese [22], disturbance, for example of guanacos [18] as well as alternance of habitat use in native species. Additionally, there might be competition with birds of prey, especially rare species such as the rufous-tailed hawk (*Buteo ventralis*) and the striated caracara (*Phalcoboenus australis*). The habitat use of feral dogs is known to vary, with some studies suggesting a preference for high-density woodlands [23], while others indicate dogs adapt to the most available habitats [10], making them potentially impact all native species. On Navarino Island, Contardo et al. [19] observed that feral dogs were frequently recorded by camera traps near landfills and farms. This finding aligns with Schüttler et al. [18], who reported that feral dogs on CHBR continue to feed on livestock. Although dogs likely consume anthropogenic food sources as part of their diet, they pose a threat to local wildlife, with the most severe impact on waterfowl populations [22] and the southernmost population of guanacos (*Lama guanicoe*), which are at risk of local extinction [24].

To assess the spread of invasive species, species distribution models (SDMs) are widely used [25, 26]. These models use occurrence records, which can include presence-absence or presence-only data [27]. However, confirming a species’ absence is often more challenging than confirming its presence [28]. As a result, SDMs based on presence-only data, such as MaxEnt, have gained popularity in recent years [29]. MaxEnt applies the principle of maximum entropy to model the relationship between presence-only data and environmental variables, helping to estimate species’ ecological niches and their potential geographic ranges [30]. Changes in population size and species diversity can lead to trophic cascades, causing potential changes in land cover [31, 32].

In this study, we investigate the landscape use of feral dogs on Navarino Island by tracking their movements and modelling habitat use. We utilise presence-only records of feral dogs and locally endangered prey species such as guanacos, upland geese (*Chloephaga picta*), flightless steamer ducks (*Tachyeres pteneres*), and two vulnerable birds of prey (*Phalcoboenus australis* and *Buteo ventralis*) to model species distribution considering prey availability, habitat structure, and predation pressure by domestic dogs, as well as to address the extent of habitat overlap as a measure for conservation needs.

## 2. Methods

### Study Area

The study was carried out on Navarino Island (see Figure 1), situated within the Cape Horn Biosphere Reserve in southern Chile. This region is part of the Magellanic sub-Antarctic ecoregion, one of the least disturbed environments on Earth, with over 90% of the area exhibiting minimal human impact [33]. The island’s landscape is dominated by southern beech forests (*Nothofagus spp*.), Magellanic tundra (*Sphagnum spp*.), shrublands, high-Andean ecosystems, and glaciers [34]. Navarino Island is sparsely populated, with approximately 2,200 residents, the majority living in Puerto Williams, the capital of the Chilean Antarctic Province. The northern coast, where fewer than ten farms are located, is the only part of the island with terrestrial infrastructure, connecting Puerto Williams to Puerto Navarino.

**Figure 1:**
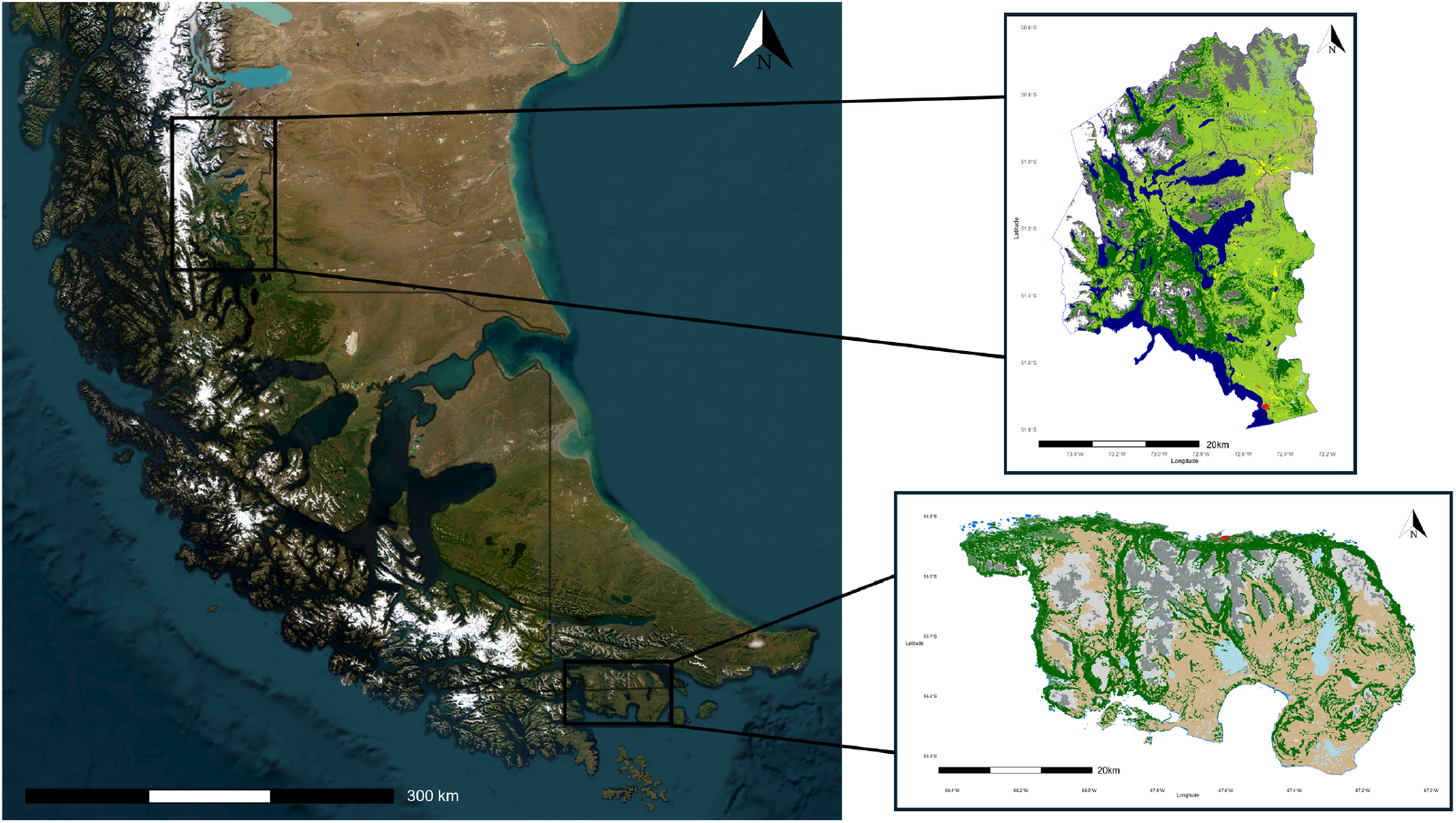
Geographical location and detailed land cover maps of study areas. Satelitte map shows the location of the two study areas, Torres del Paine National Park and Navarino Island, within the respective Chilean administrative and regional boundaries. Study areas are marked with a bounding box at their respective location and illustrated as a zoom plot with their land cover. The North arrow at the top right indicates the geographical direction, the scale at the bottom left indicates size ratio of the map.

The domestic dog population in Puerto Williams is estimated at 1.4 dogs per household, with 30.6% of these dogs reported as being unconfined by their owners [18]. However, seasonal street censuses suggest a higher number of dogs, ranging from 125 to 160, the majority of which have owners or caregivers and are not entirely abandoned. There is also considerable evidence of a feral dog population on the island. Local inhabitants frequently report sightings of suspected feral dogs, with sightings as far as 19.4 km from the northern inhabited coastline [18]. Camera traps have captured images of dogs not identifiable nor recognised as belonging to any known farms or owners [19].

Additionally, we selected Torres del Paine as a comparision site, due to comparable size (approx. 2,400 km^2^ for Torres del Paine and 2514 km^2^ for Navarino Island). Torres del Paine National Park is one of the eleven protected areas in the Magallanes Region and Chilean Antarctica. Among the most common mammals in the park are guanacos (*Lama guanicoe*). Other notable mammals include the South American grey fox (*Lycalopex griseus*), pumas (*Puma concolor*), and the endangered Chilean huemul (*Hippocamelus bisulcus*). The park supports breeding populations of 15 raptor species. These include the Andean condor (*Vultur gryphus*), black-chested buzzard-eagle (*Geranoaetus melanoleucus*), the vulnerable rufous-tailed hawk (*Buteo ventralis*), cinereous harrier (*Circus cinereus*), chimango caracara (*Milvago chimango*), crested caracara (*Caracara plancus*), Magellanic horned owl (*Bubo magellanicus*), and the austral pygmy-owl (*Glaucidium nanum*) [35]. In Torres del Paine, there have been no records of invasive feral dogs, minks or beavers to date. Free-ranging domestic horses and domestic cattle are present in two areas of the National Park, but no records of invasive predators such as minks, rats or feral dogs exist. To depict the impact of invasive predators, we decided to select a highly touristic area (approx. 250.000 visitors per year) to model human disturbance vs. invasive predator impacts on endemic species.

### Sampling methods

Our data was sampled during two field expeditions to Navarino Island (CHBR) from November 2022 to January 2023 and November 2023 to January 2024. During these trips, we traversed the island extensively, staying off accessible paths to record data on the focal species. Each tour lasted between 5 and 13 days. We recorded GPS coordinates for all our samples using a Garmin Montana 700i. Various presence data were collected. For feral dogs, we recorded footprints (including measurements), catalogued scat samples, and logged sightings with estimated distances. For guanacos, we documented fresh trails, dung piles, calls with estimated distances, and sightings. For striated caracaras, rufous-tailed hawks, and waterfowl, we recorded sightings only. Additionally, data from two expeditionists with recorded sightings from 2020 to 2023 were included.

In Torres del Paine, we modelled effects of climate change (NDVI changes from 2014-2024) as well as tourism numbers from 2019 - 2024 for each month of the presence of guanacos, upland geese as well as ashy-headed geese (*Chloephaga poliocephala*). Data were obtained from regular monitoring of CONAF within Torres del Paine National Park from 2019-2024. For 2019, 2020 and 2022, there was no data on guanacos recorded due to the Covid-19 pandemic. In Torres del Paine, guanacos were counted using fixed transects (n=13, out of which 10 are monitored on foot and three via vehicle with a constant speed of ¡20km/h) for species monitoring. If guanacos were observed in other territories, their presence was also recorded and added to their total abundance. The individuals were monitored by total number and categorised by age, sex and behaviour, to account for differences in habitat use. All individuals that were observed via visual or auditory observation were geo-referenced with GPS coordinates. Waterfowl were monitored twice per year together with neo-tropical waterfowl (CNAA), in summer and winter, using fixed transects (n = 27). Each transect is observed for a 20-minute period. Abundance as well as species are recorded, and then categorised by sex, behaviour and age.

### Data analysis

To ensure sample independence, we only included sightings within a 1 km radius per year for modeling. As most feral dog samples consisted of footprints, we excluded footprints with similar patterns (i.e., size) within a 2 km radius. Using data on GPS-tagged free-ranging dogs’ movement patterns on Navarino Island [36], we calculated spatial independence of free-ranging owned dogs and feral dog presence with a Mark Correlation Function (MCF). Habitat overlap between dog and guanaco presence records was calculated using Python version 3.12.3 with SciPy, pandas, and osgeo. We used MaxEnt version 3.4.3 in R Studio (version 4.3.1, package ‘dismo’) to model the distribution of feral dogs and guanacos. Environmental variables were selected for their importance to the target species’ habitat use. These included a Normalised Difference Vegetation Index (NDVI) with a 16-day interval MODIS/Terra NDVI image (250 m resolution) obtained from Google Earth Engine, a Global Digital Elevation Map (GDEM) from EarthData, and a land cover file, as well as prey availability for the feral dog model and dog presence records for the guanaco model (DOG). Since dogs are generalists and can survive in extreme environments, climatic variables were omitted from the feral dog model. A correlation analysis was conducted among environmental variables, and variables with a high correlation coefficient (*r* > 0.7) were discarded [30]. Different models were compared using ENMTools [37], varying some of the algorithm parameters including the regularisation parameter [38] along with the environmental variable functions. We compared two feature class sets using the default “autofeatures” option, and a set allowing only linear, quadratic, and product (LQP) features. The best-fitting model was selected and constructed via 10-fold cross-validation.

To calculate habitat connectivity and potential habitat use of feral dogs, we used Circuitscape 5.0 [39] in Julia (version 1.10.4). Connectivity and resistance were calculated based on feral dog occurrence records, determining a core area under the assumption that feral dogs use the entire island for movement and connecting this core area with foraging grounds or movement corridors. We selected areas with high resistance for Dientes de Navarino, the Puerto Williams centre, large water bodies, and islets with no or regularly flooded land bridges.

We evaluated the impacts of feral dog presence on upland geese, flightless steamer ducks, and guanacos using lasso regression (package ‘glmnet’) for variable reduction [40] (see Appendix) and generalised linear models (GLM) (package ‘glmmTMB’, [41]) as well as non-linear models (package ‘nls’) in R Studio. The response variables were presence-absence records and coordinates for the focal species. For dogs, guanacos and flightless steamer ducks, actual presence values were used; for upland geese, presence counts were used to model presence in suitable habitats across the island, where no data could be recorded. Absence values were used based on actual and modelled absence values and combined with presence data as well as environmental variables for modelling (n = 35260). Binomial error structure and logit link functions were fitted to the generalised linear models. Explanatory variables included MaxEnt predicted habitat suitability for dogs (DOGVALUE), log-transformed DOG presence records, NDVI, GDEM, landcover data (HABITAT), and climatic data (CLIMATE) from WorldClim. Detailed descriptions of response and predictor variables are in Table 2.

**Table 1:** Species occurrence models on Navarino Island.

| Candidate Model Sets | Response Variable | Predictor Variables |
| --- | --- | --- |
| Guanaco | OCCURRENCE | DOG + DOG.VALUE + NDVI<br>+ GDEM + HABITAT + CLIMATE |
| Chloephaga | OCCURRENCE | DOG + DOG.VALUE + NDVI<br>+ GDEM + HABITAT + CLIMATE |
| Tachyeres | OCCURRENCE | DOG + DOG.VALUE + NDVI<br>+ GDEM + HABITAT + CLIMATE |

**Table 2:**
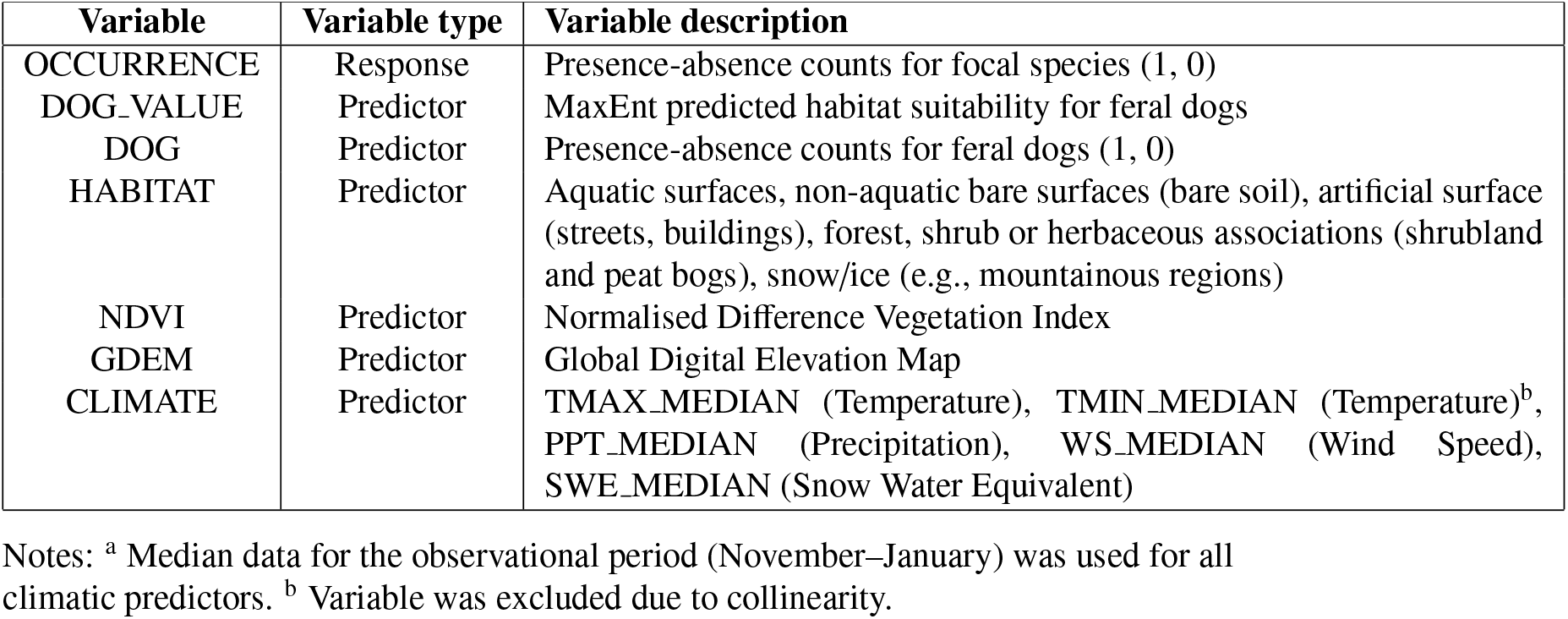
Detailed variable descriptions for GLM and NLS models.

| Variable | Variable type | Variable description |
| --- | --- | --- |
| OCCURRENCE | Response | Presence-absence counts for focal species (1, 0) |
| DOG.VALUE | Predictor | MaxEnt predicted habitat suitability for feral dogs |
| DOG | Predictor | Presence-absence counts for feral dogs (1, 0) |
| HABITAT | Predictor | Aquatic surfaces, non-aquatic bare surfaces (bare soil), artificial surface (streets, buildings), forest, shrub or herbaceous associations (shrubland and peat bogs), snow/ice (e.g., mountainous regions) |
| NDVI | Predictor | Normalised Difference Vegetation Index |
| GDEM | Predictor | Global Digital Elevation Map |
| CLIMATE | Predictor | TMAX_MEDIAN (Temperature), TMIN_MEDIAN (Temperature)<sup>b</sup>, PPT_MEDIAN (Precipitation), WS_MEDIAN (Wind Speed), SWE_MEDIAN (Snow Water Equivalent) |

Notes: <sup>a</sup> Median data for the observational period (November–January) was used for all climatic predictors. <sup>b</sup> Variable was excluded due to collinearity.

For Torres del Paine, NDVI data with a 30m resolution as well as land cover data (Sentinel-1 with 10m resolution) was obtained from Google Earth Engine. Habitat preferences as well as population stability for monitored species in Torres del Paine were calculated using Generalised Linear Models (GLMs) as well as Multinomial logistic regression (MLR) with package ‘nnet’ in R Studio. The NDVI change point analysis was conducted using the same approach as for Navarino Island.

To address the potential impacts of species decline on the ecosystem of Navarino Island, we modelled NDVI changes from 2000 to 2023 —a period during which significant ecosystem alterations have occurred on Navarino Island due to invasive species, the decline of endemic species, as well as increasing tourism and global climate change.

To capture future NDVI trends, we performed an Autoregressive Moving Average Model(ARIMA).

The ARIMA model has autoregressive (AR) terms, differencing (I), and moving average (MA) components, as well as seasonal AR, MA, and differencing terms.

The fitted model:

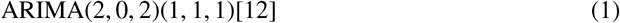

We included changes in NDVI on Navarino Island over all years in a linear model as a response and invasive species numbers, endemic species numbers, mean annual temperature, mean annual precipitation, mean annual wind speed, and an estimate of tourism increase using package ‘segmented’. We computed NDVI changes in Torres del Paine for comparison with Navarino Island using climate data from NASA with package ‘nasapower’. Data on monthly tourism numbers for Navarino Island and Torres del Paine were obtained from Servicio Nacional de Turismo (‘Sernatur’).

## 3. Results

We recorded 1261 species occurrences of striated caracaras (*n* = 2), rufous-tailed hawks (*n* = 4), flightless steamer ducks (*n* = 169), guanacos (*n* = 46), feral dogs (*n* = 476), and upland geese (*n* = 564) using all-occurrence countings (see Figure 2). Feral dogs were recorded approximately 0.19 times per square kilometer in a study area of 2514 km^2^.

**Figure 2:**
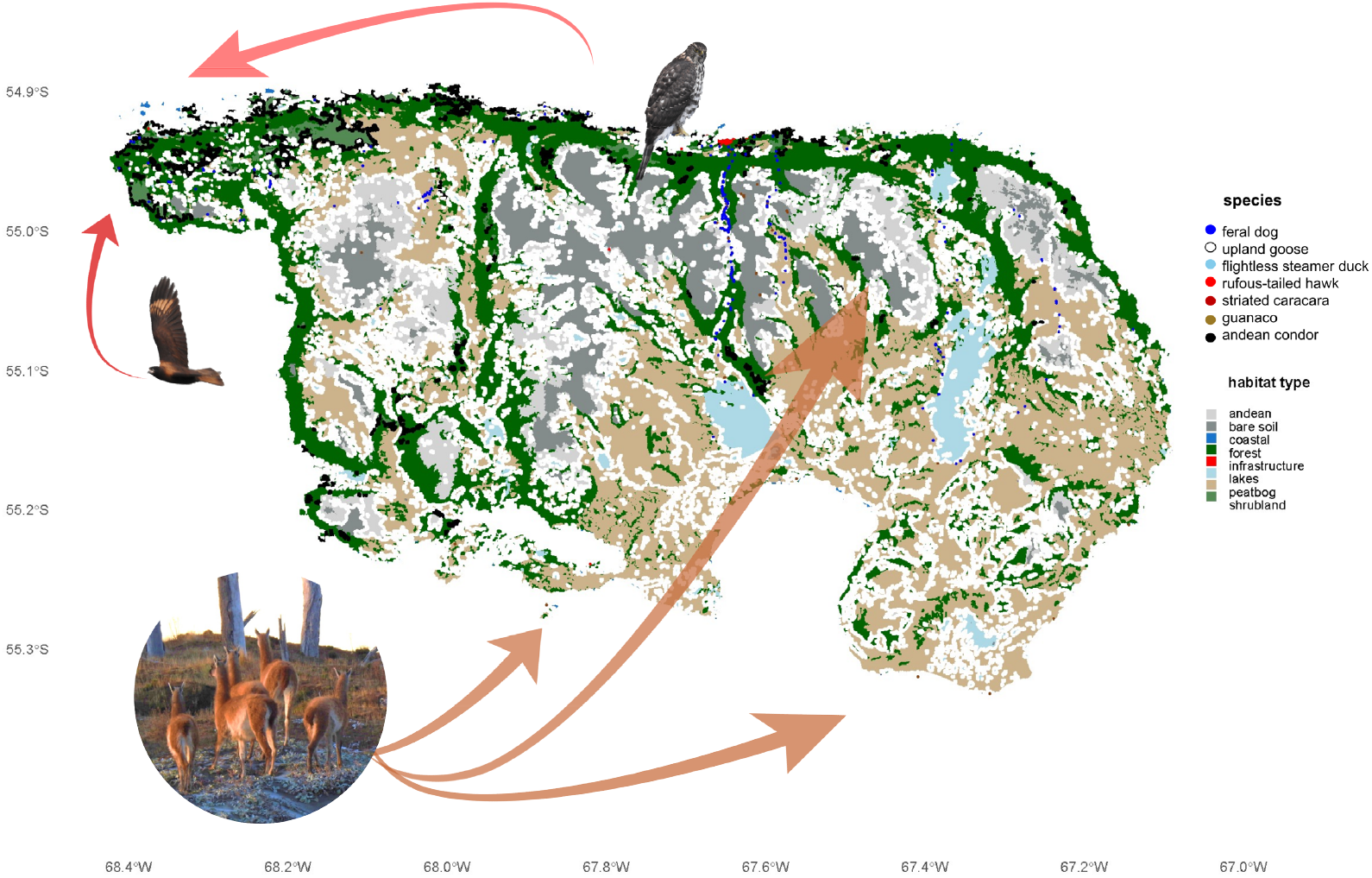
Distribution and occurrence map of all focal species recordings. X-axis and y-axis represent latitude and longitude. Coloured dots represent respective species and their associated presence records. Coloured areas indicate habitat and land cover type. Images depict respective focal species. Arrows indicate core presence areas of species.

Feral dog footprints and sightings were found to be moderately to highly independent, depending on the area. Areas closer to the North Coast, where most of the free-ranging dogs live, showed lower independence. Areas closer to the North Coast, where most of the free-ranging dogs live, showed lower independence. We generally saw repeated movement patterns of feral dogs further away from human-used paths, mostly ranging through dense forest areas. Open-area footprints and faecal samples were mostly found in further distance to the North Coast. We found a distinguishable apparence of feral dog faeces and village dog faeces, with feral dog faeces consisting of visibly more bone matter, claws, hair, and often also blood.

The predictive performance of MaxEnt models for feral dogs (AUC = 0.862) and guanacos (AUC = 0.771) was reasonably good, considering the closed setting and the limited amount of available presence data. According to the MaxEnt predictions, feral dogs prefer habitats with high NDVI and lower elevation and slope (GDEM), such as forests, shrublands, and coastal areas, as well as peat bogs (see Figure 3). Prey availability and land cover did not contribute significantly to the model’s predictive accuracy (see Table 3).

**Table 3:** Variable Contribution and Importance to MaxEnt feral dog prediction.

| Variable | Percentage of Contribution | Permutation Importance |
| --- | --- | --- |
| NDVI | 39.9 | 42.6 |
| GDEM | 38.4 | 33.6 |
| Land Cover | 16.5 | 15.6 |
| Prey Availability | 5.2 | 8.2 |

**Figure 3:**
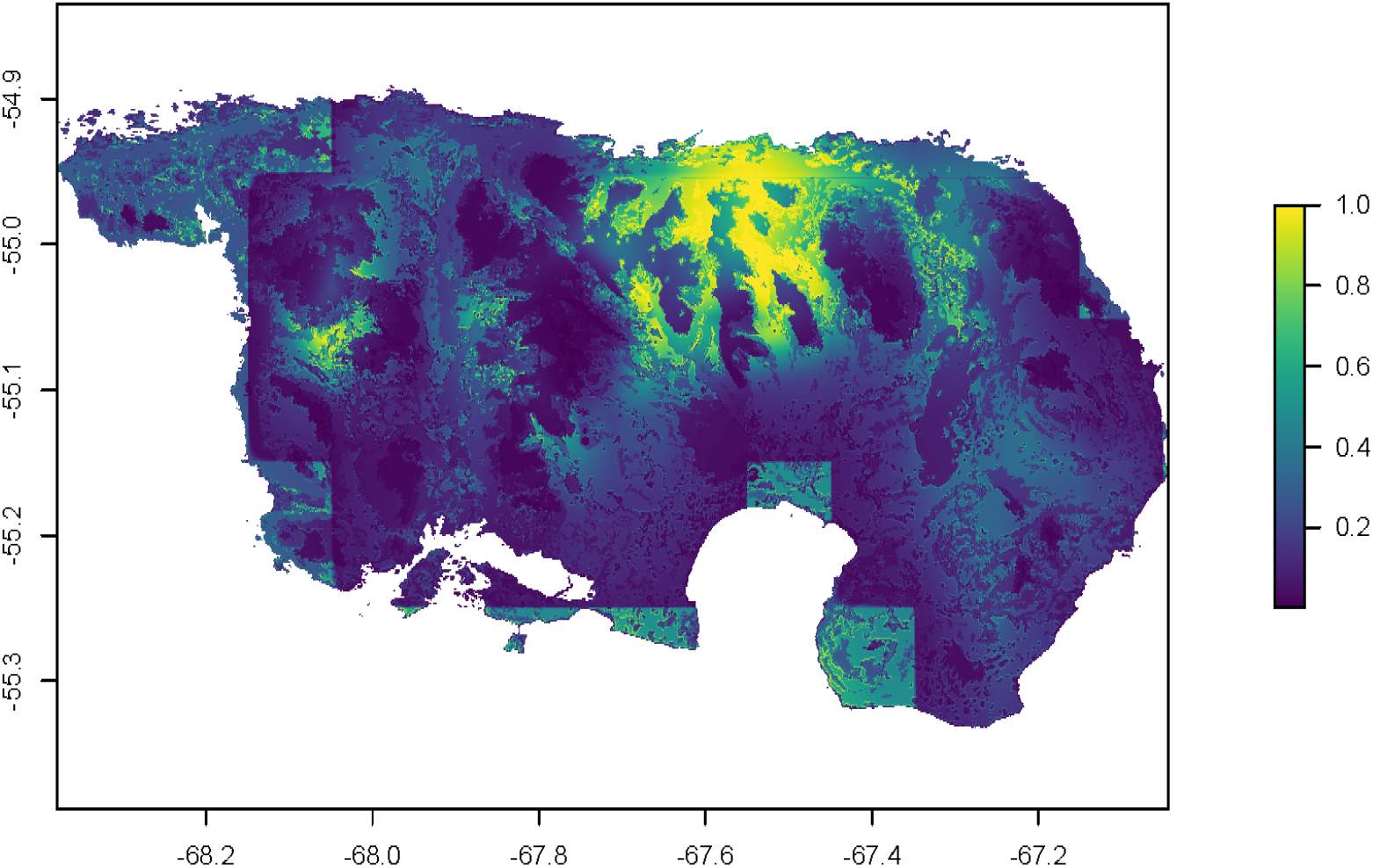
MaxEnt predicted habitat suitability for feral dogs on Navarino Island. Y-axis and X-axis indicate latitude and longitude. Gradient scale indicates habitat suitability, with values closer to zero indicating low predicted suitability and values closer to 1 indicating higher suitability.

Guanacos showed moderate habitat overlap (38.24%) with feral dogs based on their recorded occurrences (see Figure 4). Habitat suitability for guanacos was highly influenced by predicted habitat suitability for dogs, accounting for almost 68% of predicted suitability, while land cover only contributed 31%. NDVI and GDEM were negligible (see Table 4). Predicted guanaco occurrence was significantly lower in areas with high MaxEnt-predicted habitat suitability for feral dogs (χ^2^= 360 (9), β = −1.233, *p* < 0.001), but not significantly affected by actual feral dog occurrences. Areas with bare soil (χ^2^= 360 (9), β = 1.279, *p* ¡ < 0.001) and areas with potential snow and ice cover (χ^2^= 360 (9), β = 1.102, *p* < 0.001), such as mountainous regions, were positively associated with higher predicted guanaco occurrences (see Table 5).

**Table 4:** Contribution and Importance to MaxEnt prediction for Guanaco.

| Variable | Percentage of Contribution | Permutation Importance |
| --- | --- | --- |
| DOG_VALUE | 67.7 | 37.8 |
| Land Cover | 30.9 | 40.0 |
| NDVI | 1.4 | 22.2 |
| GDEM | 0.1 | 0.0 |

**Table 5:** Generalised linear model results for guanaco likelihood of occurrence.

| Variable | Estimate | Std. Error | p-value |
| --- | --- | --- | --- |
| Intercept | -2.914 | 0.133 | < 0.001 |
| DOG_VALUE |  | 0.124 | < 0.001 |
| DOG | 0.122 | 0.256 | 0.63 |
| NDVI | -0.0008 | 0.025 | 0.97 |
| GDEM | 0.0068 | 0.036 | 0.85 |
| HabitatArtificial Surface | -7.47 | 139.30 | 0.95 |
| HabitatForest | 0.414 | 0.138 | 0.002 |
| HabitatNon-Aquatic Bare Surfaces | 1.279 | 0.149 | < 0.001 |
| HabitatShrub or Herbaceous Associations | 0.354 | 0.137 | 0.009 |
| HabitatSnow / Ice | 1.102 | 0.145 | < 0.001 |

**Figure 4:**
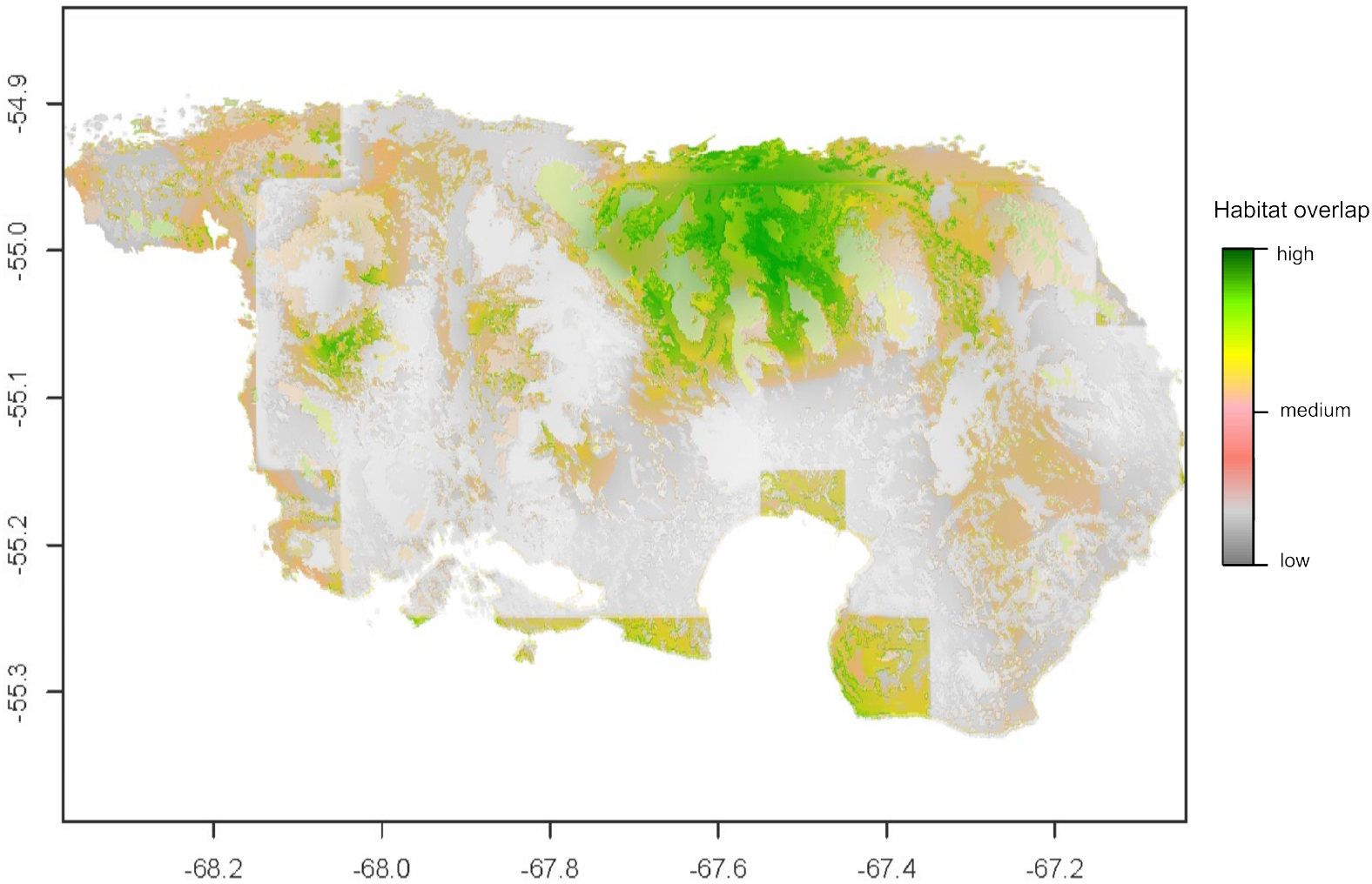
Habitat overlap of feral dogs and guanacos. Y-axis and X-axis represent latitude and longitude. Gradient scale indicates percentage of overlap, with grey depicting low overlap and green indicating high overlap.

Connectivity and resistance analysis revealed two separated areas of the island (see Figure 5). The centre of the island exhibited high connectivity with the west and south, while high resistance was found to the east. There is potentially high connectivity from the centre to the south and northwest, with restricted movement corridors due to the mountain range of the Dientes de Navarino.

**Figure 5:**
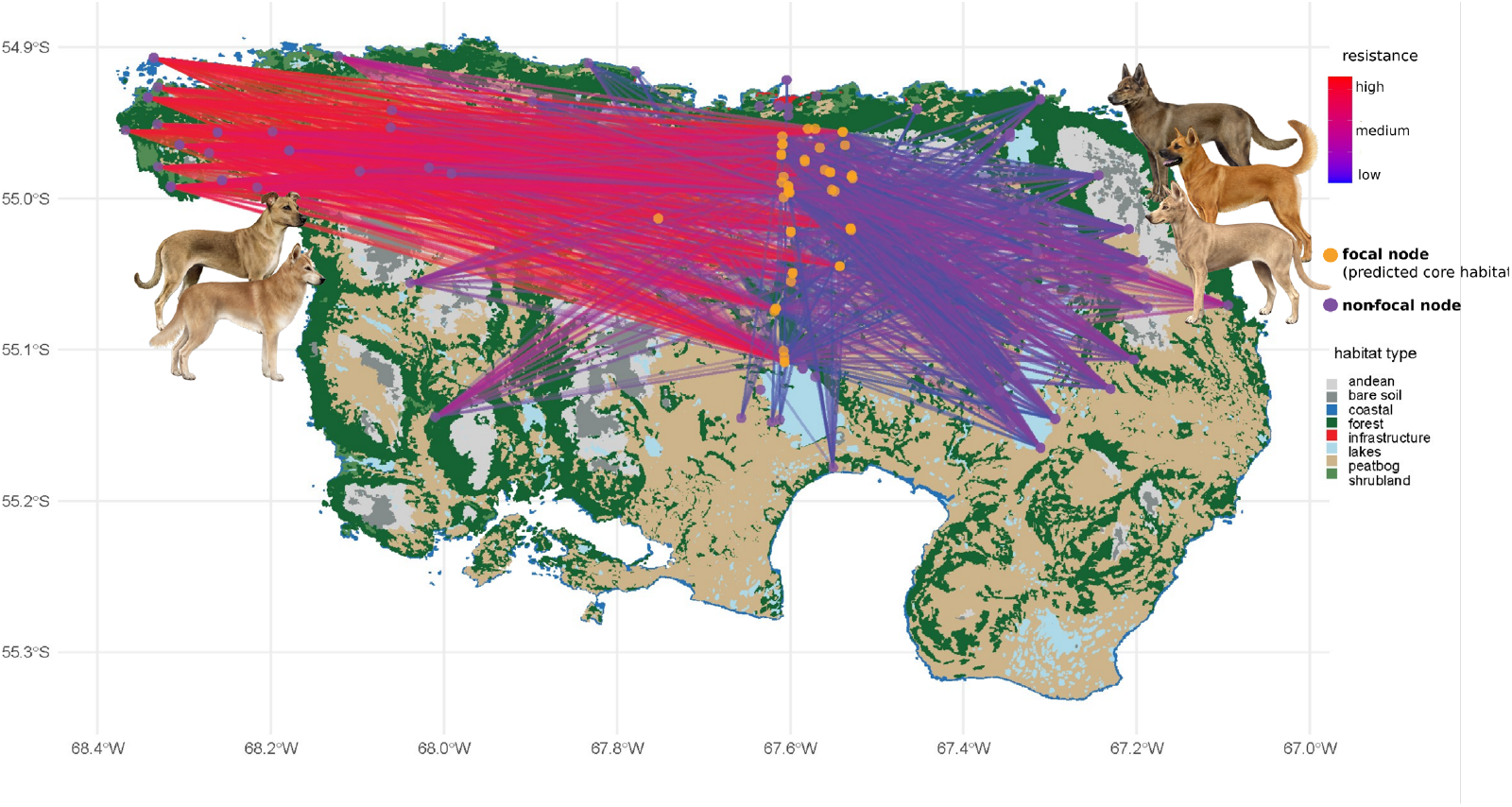
Habitat connectivity and resistance on Navarino Island. X-axis and y-axis represent latitude and longitude. Coloured dots represent focal vs. non-focal nodes. Coloured areas indicate habitat and land cover type. Gradient indicates resistance, with blue colour representing low resistance and red colour representing high resistance. Icons of dogs represent two possible distinct populations of feral dogs due to low connectivity between the West and East coast of the island.

In upland geese, lasso regression reduced the explanatory variables for occurrence modelling to eight variables, five of which were habitat types (see Appendix Table 1). These were included in a subsequent non-linear model. Upland geese showed higher predicted occurrences in forests, shrublands, and peat bogs (shrub or herbaceous associations, *p* < 0.05). Feral dog occurrences (β = −0.251, *p* < 0.001) and habitat suitability for feral dogs (β = −0.224, *p* < 0.001) were significantly negatively associated with predicted upland geese occurrence (*R*^2^ = 0.76, *p* < 0.001) (see Table 6).

**Table 6:** Non-linear model results for *Chloephaga picta* likelihood of occurrence.

| Variable | Estimate | Std. Error | p-value |
| --- | --- | --- | --- |
| Intercept | 0.5734 | 0.0784 | < 0.001 |
| DOG_VALUE | -0.252 | 0.0249 | < 0.001 |
| DOG | -0.225 | 0.0375 | < 0.001 |
| TMAX_MEDIAN | 0.0024 | 0.0078 | 0.7 |
| HABITAT | -0.0318 | 0.0031 | < 0.001 |

Controversly, the predicted occurrences of flightless steamer ducks showed no significant relationship to feral dog predicted or actual presence (*p* > 0.1). According to our GLM results (see Table 7), flightless steamer ducks were more likely to occur in areas with lower median precipitation (X^2^ = 51.03 (11), ß = −0.495, *p* < 0.001) and higher median wind speed (X^2^ = 51.03 (11), ß = 2.289, *p* < 0.001). All variables included in the GLM were also kept during variable reduction with lasso (see Appendix Table 2).

**Table 7:** Generalised linear model results for *Tachyeres pteneres* likeliness of occurrence.

| Variable | Estimate | Std. Error | z-value | p-value |
| --- | --- | --- | --- | --- |
| Intercept | 4.0300 | 4.7750 | 0.844 | 0.39 |
| DOG | 0.6157 | 1.0680 | 0.577 | 0.56 |
| DOG_VALUE | -0.5841 | 0.6054 | -0.965 | 0.33 |
| NDVI | 0.0000 | 0.0000 | 0.609 | 0.54 |
| HabitatArtificial Surface | -14.6600 | 7604.0000 | -0.002 | 0.99 |
| HabitatForest | -0.5860 | 0.5001 | -1.172 | 0.24 |
| HabitatMountain | -2.3880 | 1.0960 | -2.179 | 0.02 |
| HabitatNon-Aquatic Bare Surfaces | -15.3300 | 579.4000 | -0.026 | 0.97 |
| HabitatShrub or herbaceous associations | -0.4922 | 0.4647 | -1.059 | 0.28 |
| TMAX_MEDIAN | -0.1301 | 0.2319 | -0.561 | 0.57 |
| PPT_MEDIAN | -0.4957 | 0.1094 | -4.531 | < 0.001 |
| WS_MEDIAN | 2.2890 | 0.4879 | 4.691 | < 0.001 |

We found several carcasses of dead cows in close proximity to farmlands (n = 48), with bite marks relating to dogs. In further distance to farmlands, no cadavers or bones of domestic free-roaming animals were found. In addition, we found several dead beavers, including a sighting of feral dogs (n = 4) eating a dead beaver in the forest of the central area of the island. Feral dogs as well as free-ranging dogs have also been previously reported to consume dead whale carcasses at the North and West coast.

At the East coast, we found a dead rufous-tailed hawk with characteristic plucked remains caused by dogs. We observed rufous-tailed hawks visiting this part of the island, as well as one striated caracara. At the West and North coast, we found dog scat with muscle shells and stones as well as small bones of shorebirds. In line with these findings, there have been reports of feral dogs chasing and killing shorebirds.

In the comparative ecosystem, Torres del Paine National Park, where there are currently no records of invasive predators, guanacos had an average total population - calculated by the sum of all observations divided by the years with more than one observation - of 432.6, with juveniles contributing 52.4 and chulengos 43.4 individuals. For geese, upland geese had 417 individuals and ashy-headed geese had 91.4. The stability rate for guanacos was estimated at β = 1.926, *p* < 0.001, indicating a strong positive trend. The calculated reproduction rate for guanacos was approximately 0.096, indicating an annual growth of around 9.6%. The stability of guanacos was strongly associated with open habitats such as steppe (*X*^*2*^=1445.5(1), β = 2.508, SE = 0.119, *z* = 21.013, *p* < 0.001) or shrubland (*X*^*2*^=1445.5(1), β = 1.385, SE = 0.125, *z* = 11.036, *p* < 0.001) (see Figure 6). This is in contrast to findings on Navarino Island, where guanacos preferred mountain habitats and avoided open habitats. Considering observations of approximately 400 guanacos in the early 2000s, which now declined to approx. 35 individuals [42], the annual exponential decline rate for the Navarino population is approximately −0.128, posing a drastic contrast to the population development in Torres del Paine.

**Figure 6:**
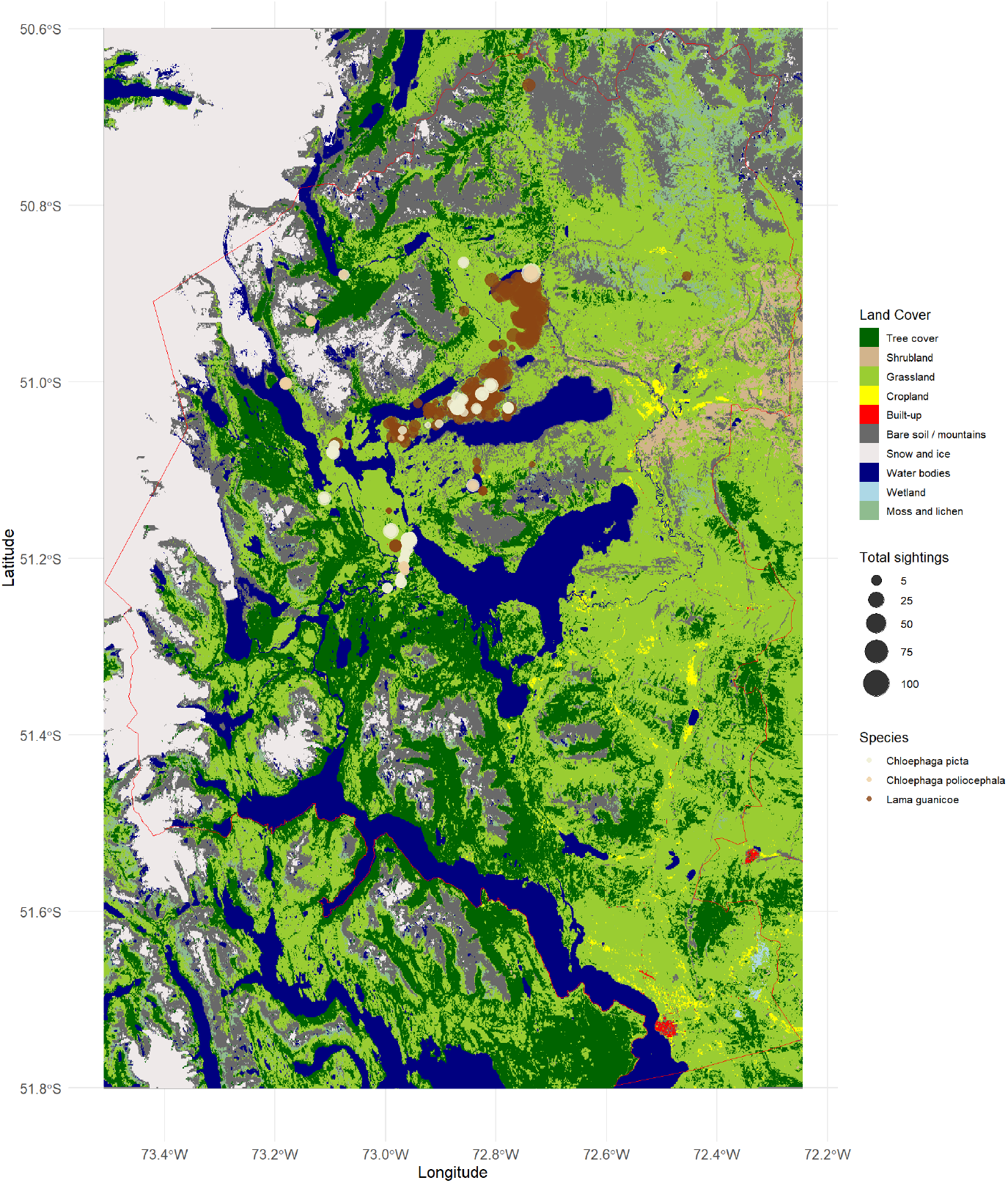
Species total observations, calculated from each individual monitoring. Coloured dots represent species, while dot size indicates total number of individuals in each observation in Torres del Paine. Red border represents boundary for Torres del Paine National Park. Colours show the land cover features in the park, obtained from Sentinel-1 land cover data from Google Earth Engine

**Figure 7:**
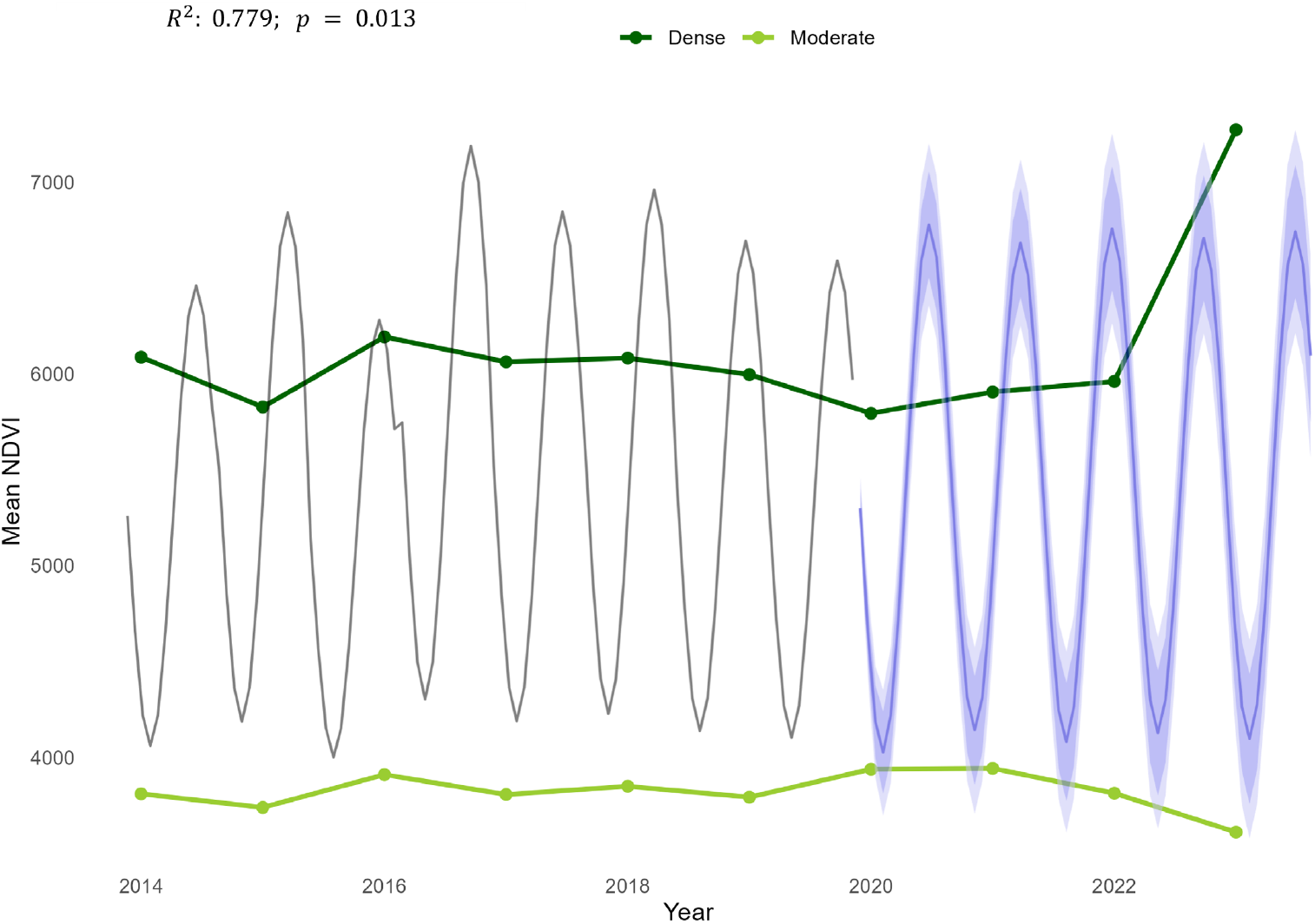
Temporal trends in mean NDVI for dense and moderate vegetation categories on Navarino Island from 2014 to 2023 with an overlay of a forecast of NDVI values using a seasonal ARIMA model. The trend lines illustrate the mean NDVI for dense (dark green) and moderate (light green) vegetation cover over time, with dense vegetation showing more variability than moderate vegetation. The black line on the left in the ARIMA represents historical NDVI data, illustrating clear seasonal fluctuations in vegetation health from 2016 to 2023. The blue line on the right shows the projected NDVI values from 2020 onward, with shaded confidence intervals capturing the prediction uncertainty.

For upland geese, the stability rate was β = 1.205, *p* = 0.045, whereas ashy-headed geese had a population stability of β = 0.892, *p* = 0.071, which, although positive, indicates a weaker stability trend compared to the other species, with more variability in population dynamics over time. Preferred growth conditions for geese were observed in wetland habitats (*G*^*2*^=18.00(4), β = 2.281, *p* = 0.066), while river habitats showed a negative but non-significant association (*G*^*2*^=18.00(4),β = −0.360, *p* = 0.706). The growth rate of *Chloephaga picta* in wetland habitats was estimated at approximately 0.00062, whereas it was slightly negative in river habitats (−0.00029). Due to a lack of data and insufficient monitoring on Navarino Island, stability rates for upland geese and flightless steamer ducks on Navarino Island cannot be provided.

### 3.1 Modelling ecosystem impacts

### 3.1.1 Navarino Island

The NDVI time series analysis from 2000-2023 revealed several notable breakpoints in the data, indicating periods of statistically significant change on Navarino Island. The breakpoints were found at 2014 (SE = 0.137), 2016 (SE = 0.076) and additionally between 2017 and 2019 (SE = 0.221; 0.183; 0.159), suggesting a high level of statistical confidence in shifts occurring around 2014 and 2016, compared to later shifts. The 1-breakpoint model, with a breakpoint at mid-2014 (BIC = 1695, RSS = 229758343), was selected as the optimal model based on the lowest Bayesian Information Criterion (BIC). The 4-breakpoint model, with the lowest residual sum of squares (RSS), identifies breakpoints at December 2014, June 2016, January 2018, and July 2019 (BIC = 1722; RSS = 216092796), thereby capturing multiple structural changes over time. Lower RSS values indicate a closer fit to the data and thus enhance model accuracy by reducing unexplained variance, allowing us to detect more subtle environmental or ecological shifts.

Although the temporal NDVI trend analysis with predictions based on data from 2000 - 2020 was closely related to the actual shifts (see Appendix) and revealed significant changes over the years in the linear model (R^2^: 0.779; *p* = 0.013), only one of the model coefficients (beaver abundance) had a statistically significant effect on these temporal shifts. Decreasing guanaco numbers showed a slight association with NDVI changes, while cows - invasive ungulates - had a negative effect on increasing NDVI, potentially associated with population growth in invasive ungulates and lower predation rates on cows and horses than on guanacos. The most pronounced changes were observed in areas of dense and moderate vegetation, with an increase in dense vegetation, such as forests, and a decrease in shrubland and bush cover (see Figure 8). In contrast to NDVI shifts on Navarino Island, the vegetation density and productivity in Torres del Paine showed an opposite temporal trend, with decreasing forest density and increasing plains. No breakpoints could be detected in Torres del Paine for NDVI trend, but an association with increasing guanaco density as well as increasing median annual temperature was found (see Table 8).

**Table 8:** Linear model coefficients for NDVI change from 2000-2023.

| Variable | Estimate | Std. Error | t-value | p-value |
| --- | --- | --- | --- | --- |
| (Intercept) | -2.787e+07 | 2.061e+07 | -1.352 | 0.213 |
| Year | -8.647e+06 | 7.797e+06 | -1.109 | 0.300 |
| Beavers | 9.411e+02 | 7.958e+02 | 1.183 | 0.027 |
| Guanacos | -3.314e+03 | 3.236e+03 | 1.024 | 0.063 |
| Cows | -1.598e+06 | 1.383e+06 | -1.155 | 0.281 |
| Dogs | -9.222e+01 | 1.419e+02 | -0.650 | 0.534 |
| Temperature | -1.848e+02 | 1.243e+03 | -0.149 | 0.885 |
| Precipitation | -2.116e+00 | 4.891e+00 | -0.433 | 0.677 |
| WindSpeed | 2.798e+01 | 3.253e+02 | 0.086 | 0.934 |
| TourismIndex | 1.112e+03 | 6.777e+02 | 1.641 | 0.140 |

**Table 9:** Lasso regression results for upland geese.

| **Variable** | **Lasso Estimate** |
| --- | --- |
| Intercept | 0.3965481389 |
| Dog | -0.0290831224 |
| Dog_Value | -0.0449296317 |
| NDVI | . |
| GDEM | . |
| tmax_median | 0.0204318225 |
| ppt_median | . |
| ws_median | . |
| HabitatAquatic Surfaces | 0.0170204009 |
| HabitatArtificial Surface | 0.0004680223 |
| HabitatForest | 0.0327422939 |
| HabitatMountain | . |
| HabitatNon-Aquatic Bare Surfaces | -0.0147092666 |
| HabitatShrub or herbaceous associations | -0.0184511664 |

**Table 10:**
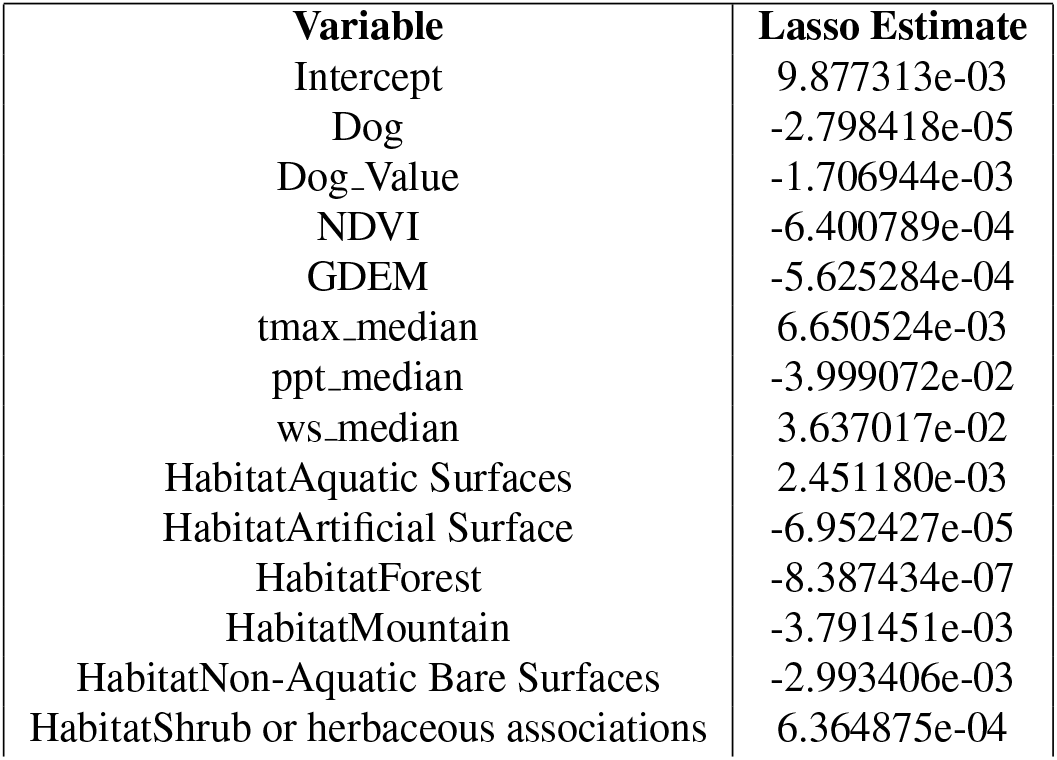
Lasso regression results for flightless steamer ducks.

| **Variable** | **Lasso Estimate** |
| --- | --- |
| Intercept | 9.877313e-03 |
| Dog | -2.798418e-05 |
| Dog_Value | -1.706944e-03 |
| NDVI | -6.400789e-04 |
| GDEM | -5.625284e-04 |
| tmax_median | 6.650524e-03 |
| ppt_median | -3.999072e-02 |
| ws_median | 3.637017e-02 |
| HabitatAquatic Surfaces | 2.451180e-03 |
| HabitatArtificial Surface | -6.952427e-05 |
| HabitatForest | -8.387434e-07 |
| HabitatMountain | -3.791451e-03 |
| HabitatNon-Aquatic Bare Surfaces | -2.993406e-03 |
| HabitatShrub or herbaceous associations | 6.364875e-04 |

**Table 11:** Segmented Model Summary.

| Measure | Min | 1st Q | Median | Mean | 3rd Q | Max |
| --- | --- | --- | --- | --- | --- | --- |
| R<sup>2</sup> | 0.0001383 | 0.0469437 | 0.0816442 | 0.0937370 | 0.1304141 | 0.6361754 |
| RMSE | 0.006739 | 0.038707 | 0.069683 | 0.073201 | 0.104296 | 0.219844 |
| AIC | -777.8 | -393.2 | -263.9 | -296.4 | -175.1 | -11.1 |

**Table 12:** Model Coefficients for the influence on NDVI. Weather date originated from [43], with T2M = temperature in 2 m height, PRECTOT-CORR = total corrected precipitation, and RH2M = relative humidity in 2 m height. The variable visitor represents tourists per month. Week is representative for time of the year to estimate seasonal influence.

Table .11: Segmented Model Summary
| Year | Term | Estimate | Std.Error | t.value | p.value | edf_value |
| --- | --- | --- | --- | --- | --- | --- |
| 2015 | (Intercept) | 0.07 | 0.00 | 129.67 | 0.00 |  |
| 2015 | s(PRECTOTCORR).1 | -0.04 | 0.00 | -15.42 |  | 2.99 |
| 2015 | s(PRECTOTCORR).2 | -0.08 | 0.01 | -13.61 |  | 2.99 |
| 2015 | s(PRECTOTCORR).3 | -0.02 | 0.00 | -4.91 |  | 2.99 |
| 2015 | s(RH2M).1 | 0.00 | 0.00 | 1.03 |  | 2.98 |
| 2015 | s(RH2M).2 | 0.06 | 0.00 | 19.09 |  | 2.98 |
| 2015 | s(RH2M).3 | 0.02 | 0.00 | 10.35 |  | 2.98 |
| 2015 | s(T2M).1 | -0.03 | 0.00 | -12.02 |  | 2.98 |
| 2015 | s(T2M).2 | 0.01 | 0.00 | 4.37 |  | 2.98 |
| 2015 | s(T2M).3 | 0.04 | 0.00 | 21.03 |  | 2.98 |
| 2015 | s(week).1 | 0.03 | 0.00 | 15.36 |  | 2.00 |
| 2015 | s(week).2 | 0.07 | 0.00 | 37.35 |  | 2.00 |
| 2016 | (Intercept) | 0.06 | 0.00 | 146.76 | 0.00 |  |
| 2016 | s(PRECTOTCORR).1 | 0.00 | 0.00 | 2.37 |  | 2.88 |
| 2016 | s(PRECTOTCORR).2 | 0.02 | 0.01 | 3.42 |  | 2.88 |
| 2016 | s(PRECTOTCORR).3 | -0.02 | 0.00 | -10.07 |  | 2.88 |
| 2016 | s(RH2M).1 | -0.03 | 0.00 | -16.14 |  | 2.99 |
| 2016 | s(RH2M).2 | 0.04 | 0.00 | 16.88 |  | 2.99 |
| 2016 | s(RH2M).3 | 0.02 | 0.00 | 14.52 |  | 2.99 |
| 2016 | s(T2M).1 | -0.03 | 0.00 | -14.88 |  | 2.99 |
| 2016 | s(T2M).2 | 0.01 | 0.00 | 4.78 |  | 2.99 |
| 2016 | s(T2M).3 | 0.04 | 0.00 | 19.75 |  | 2.99 |
| 2016 | s(week).1 | 0.04 | 0.00 | 42.09 |  | 2.00 |
| 2016 | s(week).2 | 0.10 | 0.00 | 52.95 |  | 2.00 |
| 2017 | (Intercept) | 0.06 | 0.00 | 161.67 | 0.00 |  |
| 2017 | s(PRECTOTCORR).1 | -0.00 | 0.00 | -2.72 |  | 2.90 |
| 2017 | s(PRECTOTCORR).2 | -0.03 | 0.01 | -6.15 |  | 2.90 |
| 2017 | s(PRECTOTCORR).3 | -0.01 | 0.00 | -6.25 |  | 2.90 |
| 2017 | s(RH2M).1 | 0.02 | 0.00 | 12.06 |  | 2.99 |
| 2017 | s(RH2M).2 | 0.07 | 0.00 | 30.99 |  | 2.99 |
| 2017 | s(RH2M).3 | 0.01 | 0.00 | 5.43 |  | 2.99 |
| 2017 | s(T2M).1 | -0.01 | 0.00 | -4.60 |  | 2.97 |
| 2017 | s(T2M).2 | -0.02 | 0.00 | -10.83 |  | 2.97 |
| 2017 | s(T2M).3 | 0.03 | 0.00 | 18.19 |  | 2.97 |
| 2017 | s(week).1 | 0.00 | 0.00 | 3.81 |  | 2.00 |
| 2017 | s(week).2 | 0.05 | 0.00 | 38.72 |  | 2.00 |
| 2018 | (Intercept) | 0.06 | 0.00 | 151.32 | 0.00 |  |
| 2018 | s(PRECTOTCORR).1 | -0.02 | 0.00 | -12.65 |  | 2.99 |
| 2018 | s(PRECTOTCORR).2 | -0.00 | 0.00 | -0.05 |  | 2.99 |
| 2018 | s(PRECTOTCORR).3 | -0.04 | 0.00 | -20.88 |  | 2.99 |
| 2018 | s(RH2M).1 | 0.00 | 0.00 | 2.85 |  | 2.99 |
| 2018 | s(RH2M).2 | 0.08 | 0.00 | 31.06 |  | 2.99 |
| 2018 | s(RH2M).3 | -0.00 | 0.00 | -1.53 |  | 2.99 |
| 2018 | s(T2M).1 | -0.02 | 0.00 | -7.96 |  | 2.98 |
| 2018 | s(T2M).2 | -0.06 | 0.00 | -21.72 |  | 2.98 |
| 2018 | s(T2M).3 | 0.04 | 0.00 | 23.07 |  | 2.98 |
| 2018 | s(week).1 | -0.00 | 0.00 | -0.46 |  | 2.00 |
| 2018 | s(week).2 | 0.11 | 0.00 | 55.57 |  | 2.00 |
| 2019 | (Intercept) | 0.06 | 0.00 | 153.60 | 0.00 |  |
| 2019 | s(PRECTOTCORR).1 | -0.00 | 0.00 | -4.96 |  | 2.96 |
| 2019 | s(PRECTOTCORR).2 | -0.04 | 0.00 | -9.60 |  | 2.96 |
| 2019 | s(PRECTOTCORR).3 | -0.02 | 0.00 | -15.49 |  | 2.96 |
| 2019 | s(RH2M).1 | -0.02 | 0.00 | -9.62 |  | 2.99 |
| 2019 | s(RH2M).2 | 0.05 | 0.00 | 27.30 |  | 2.99 |
| 2019 | s(RH2M).3 | -0.01 | 0.00 | -3.81 |  | 2.99 |
| 2019 | s(T2M).1 | -0.02 | 0.00 | -12.26 |  | 2.99 |
| 2019 | s(T2M).2 | -0.01 | 0.00 | -4.89 |  | 2.99 |
| 2019 | s(T2M).3 | 0.06 | 0.00 | 40.76 |  | 2.99 |
| 2019 | s(visitors).1 | 0.32 | 0.01 | 48.17 |  | 3.00 |
| 2019 | s(visitors).2 | 0.21 | 0.00 | 45.81 |  | 3.00 |
| 2019 | s(visitors).3 | 0.18 | 0.00 | 46.32 |  | 3.00 |
| 2019 | s(week).1 | 0.03 | 0.00 | 22.11 |  | 2.00 |
| 2019 | s(week).2 | 0.12 | 0.00 | 33.85 |  | 2.00 |
| 2020 | (Intercept) | 0.05 | 0.00 | 152.20 | 0.00 |  |
| 2020 | s(PRECTOTCORR).1 | -0.02 | 0.00 | -13.90 |  | 2.99 |
| 2020 | s(PRECTOTCORR).2 | 0.01 | 0.00 | 2.99 |  | 2.99 |
| 2020 | s(PRECTOTCORR).3 | -0.01 | 0.00 | -5.54 |  | 2.99 |
| 2020 | s(RH2M).1 | 0.04 | 0.00 | 27.22 |  | 3.00 |
| 2020 | s(RH2M).2 | 0.07 | 0.00 | 33.21 |  | 3.00 |
| 2020 | s(RH2M).3 | 0.04 | 0.00 | 24.54 |  | 3.00 |
| 2020 | s(T2M).1 | 0.03 | 0.00 | 13.02 |  | 2.99 |
| 2020 | s(T2M).2 | -0.01 | 0.00 | -3.94 |  | 2.99 |
| 2020 | s(T2M).3 | 0.10 | 0.00 | 51.43 |  | 2.99 |
| 2020 | s(visitors).1 | -0.01 | 0.06 | -0.10 |  | 2.05 |
| 2020 | s(visitors).2 | -0.04 | 0.02 | -1.73 |  | 2.05 |
| 2020 | s(visitors).3 | -0.01 | 0.01 | -0.50 |  | 2.05 |
| 2020 | s(week).1 | 0.03 | 0.00 | 29.67 |  | 2.00 |
| 2020 | s(week).2 | 0.12 | 0.00 | 48.39 |  | 2.00 |
| 2021 | (Intercept) | 0.05 | 0.00 | 138.28 | 0.00 |  |
| 2021 | s(PRECTOTCORR).1 | -0.04 | 0.00 | -30.55 |  | 3.00 |
| 2021 | s(PRECTOTCORR).2 | -0.09 | 0.00 | -25.69 |  | 3.00 |
| 2021 | s(PRECTOTCORR).3 | 0.03 | 0.00 | 15.44 |  | 3.00 |
| 2021 | s(RH2M).1 | 0.01 | 0.00 | 8.99 |  | 2.99 |
| 2021 | s(RH2M).2 | 0.07 | 0.00 | 36.92 |  | 2.99 |
| 2021 | s(RH2M).3 | -0.00 | 0.00 | -1.56 |  | 2.99 |
| 2021 | s(T2M).1 | -0.00 | 0.00 | -2.16 |  | 2.91 |
| 2021 | s(T2M).2 | -0.01 | 0.00 | -6.44 |  | 2.91 |
| 2021 | s(T2M).3 | 0.05 | 0.00 | 35.90 |  | 2.91 |
| 2021 | s(visitors).1 | 0.04 | 0.00 | 23.16 |  | 3.00 |
| 2021 | s(visitors).2 | -0.03 | 0.00 | -19.38 |  | 3.00 |
| 2021 | s(visitors).3 | 0.05 | 0.00 | 28.45 |  | 3.00 |
| 2021 | s(week).1 | 0.06 | 0.00 | 57.69 |  | 2.00 |
| 2021 | s(week).2 | 0.09 | 0.00 | 83.27 |  | 2.00 |
| 2022 | (Intercept) | 0.05 | 0.00 | 153.23 | 0.00 |  |
| 2022 | s(PRECTOTCORR).1 | 0.01 | 0.00 | 12.34 |  | 2.98 |
| 2022 | s(PRECTOTCORR).2 | 0.02 | 0.00 | 8.00 |  | 2.98 |
| 2022 | s(PRECTOTCORR).3 | -0.04 | 0.00 | -28.69 |  | 2.98 |
| 2022 | s(RH2M).1 | -0.02 | 0.00 | -13.55 |  | 2.99 |
| 2022 | s(RH2M).2 | 0.03 | 0.00 | 24.37 |  | 2.99 |
| 2022 | s(RH2M).3 | 0.02 | 0.00 | 14.95 |  | 2.99 |
| 2022 | s(T2M).1 | -0.01 | 0.00 | -6.55 |  | 2.95 |
| 2022 | s(T2M).2 | -0.02 | 0.00 | -9.11 |  | 2.95 |
| 2022 | s(T2M).3 | 0.03 | 0.00 | 31.01 |  | 2.95 |
| 2022 | s(visitors).1 | -0.14 | 0.01 | -24.18 |  | 3.00 |
| 2022 | s(visitors).2 | -0.03 | 0.00 | -25.33 |  | 3.00 |
| 2022 | s(visitors).3 | -0.14 | 0.01 | -21.37 |  | 3.00 |
| 2022 | s(week).1 | -0.05 | 0.00 | -19.59 |  | 2.00 |
| 2022 | s(week).2 | 0.03 | 0.00 | 24.34 |  | 2.00 |
| 2023 | (Intercept) | 0.04 | 0.00 | 188.80 | 0.00 |  |
| 2023 | s(PRECTOTCORR).1 | 0.01 | 0.00 | 15.32 |  | 2.99 |
| 2023 | s(PRECTOTCORR).2 | -0.02 | 0.00 | -8.08 |  | 2.99 |
| 2023 | s(PRECTOTCORR).3 | 0.00 | 0.00 | 1.20 |  | 2.99 |
| 2023 | s(RH2M).1 | 0.00 | 0.00 | 3.67 |  | 2.99 |
| 2023 | s(RH2M).2 | 0.05 | 0.00 | 35.21 |  | 2.99 |
| 2023 | s(RH2M).3 | 0.00 | 0.00 | 2.35 |  | 2.99 |
| 2023 | s(T2M).1 | 0.01 | 0.00 | 8.67 |  | 2.99 |
| 2023 | s(T2M).2 | -0.04 | 0.00 | -26.53 |  | 2.99 |
| 2023 | s(T2M).3 | 0.06 | 0.00 | 52.52 |  | 2.99 |
| 2023 | s(visitors).1 | -0.02 | 0.00 | -3.16 |  | 2.99 |
| 2023 | s(visitors).2 | 0.09 | 0.00 | 27.33 |  | 2.99 |
| 2023 | s(visitors).3 | -0.02 | 0.00 | -5.73 |  | 2.99 |
| 2023 | s(week).1 | 0.03 | 0.00 | 39.02 |  | 2.00 |
| 2023 | s(week).2 | 0.13 | 0.00 | 59.97 |  | 2.00 |

**Figure 8:**
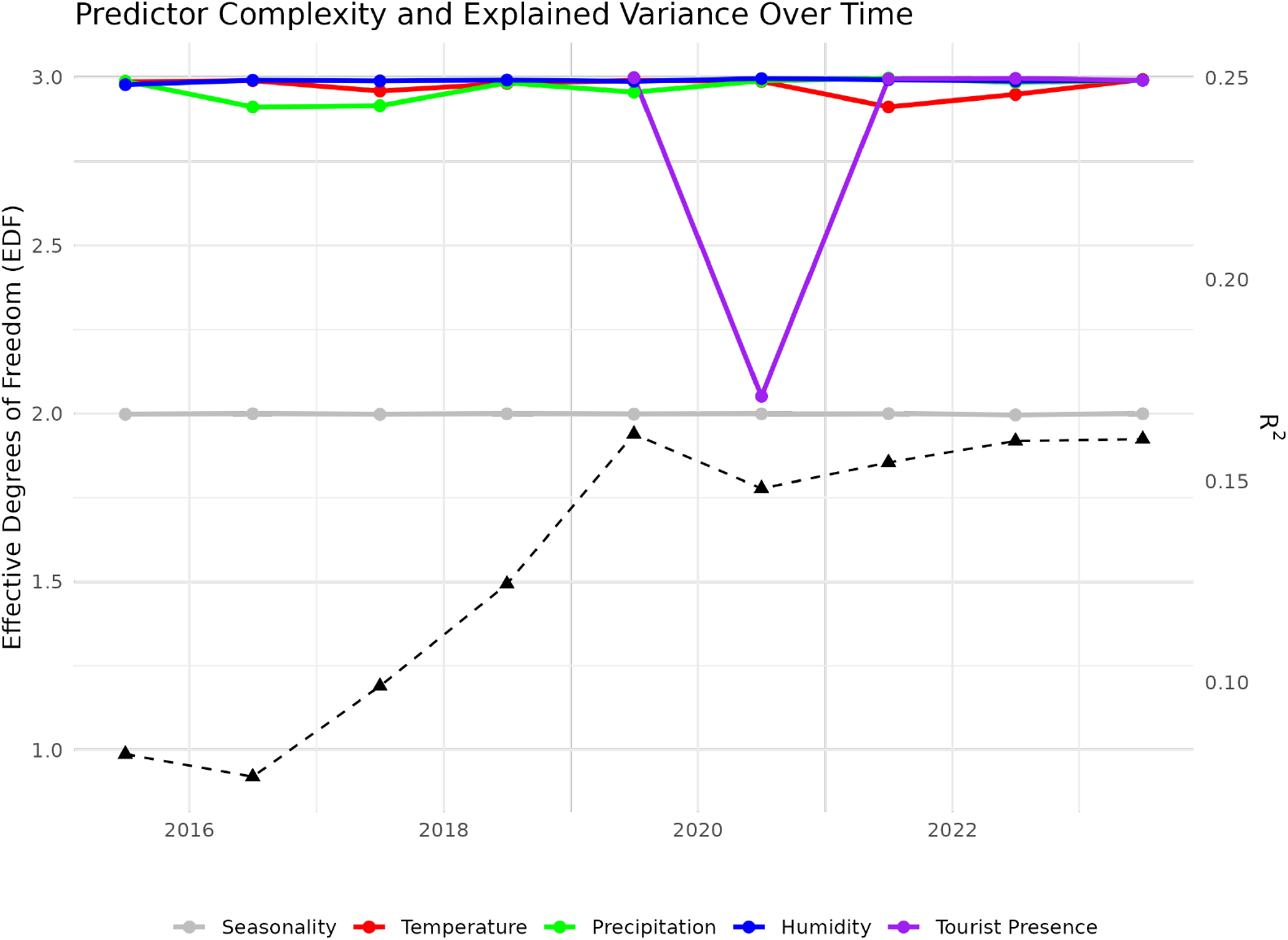
Predictor Complexity and Explained Variance of NDVI in Torres del Paine calculated separately for each year from 2015 to 2023. Colored lines show the climatic variables (temperature, corrected precipitation and relative humidity (based on [43]), and tourist presence, gray line indicates a constant influence of seasonality represented by weeks (left y-axis). Dashed black line shows the R^2^ for each year (right y-axis).

**Figure 9:**
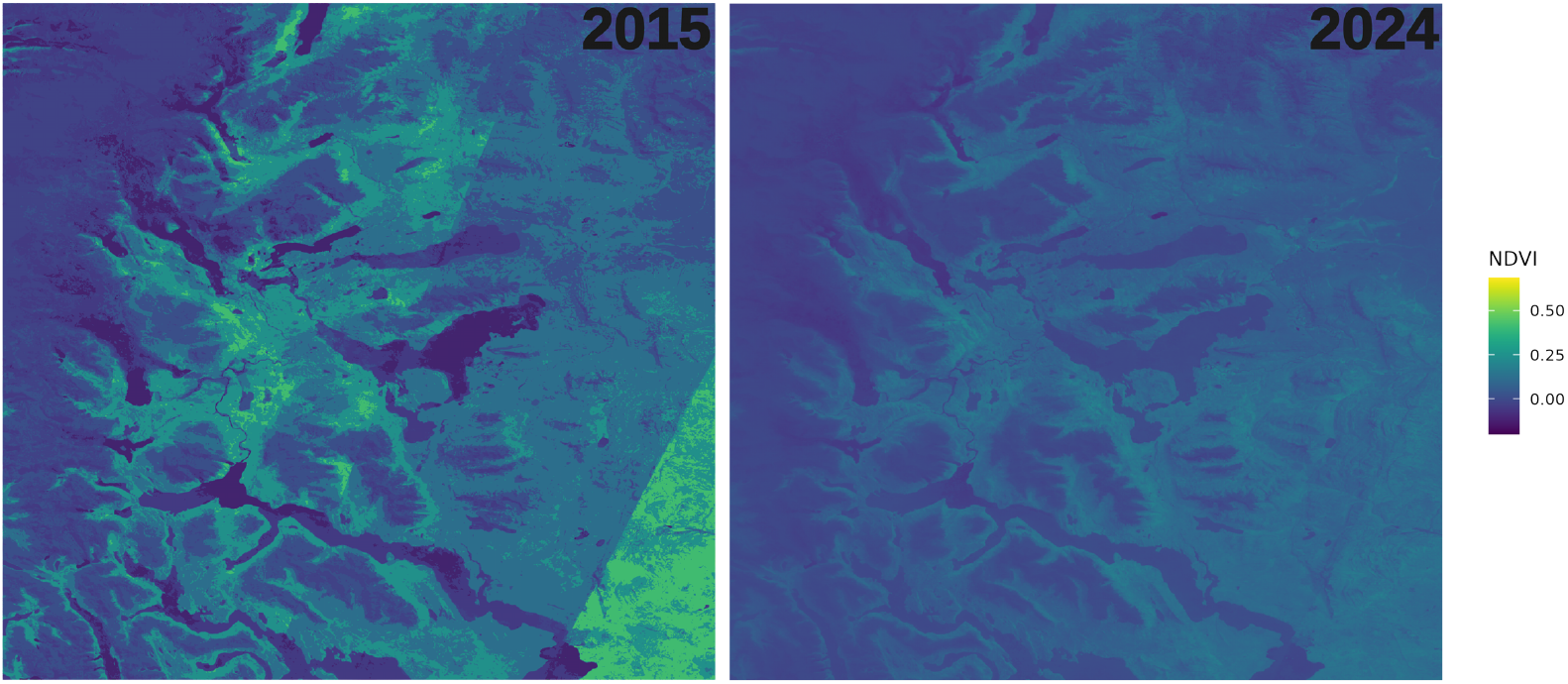
NDVI for the beginning of 2015 (left) and 2024 (right). This indicates a shift towards less vegetation and water.

#### 3.1.2 Torres del Paine

We obtained 1.5 million segmented models for the NDVI of Torres del Paine based on each area (by longitude and latitude) for each year. The R^2^ (median 0.087), RSME (median 0.069), and AIC (median -363) of these models indicate a low but constant estimation (see appendix). Thus, the location and time are not the main influences on NDVI. From 2015 to 2023 it is influenced in a non-linear relation of temperature, precipitation, and humidity of similar strength 8. Seasonality over the years remains, as expected, constant and non-linear on a lower level. The number of tourists (which is only available from 2019 on) has a non-linear effect as strong as the climatic variables, except for the years when travelling was reduced due to the COVID-19 pandemic. It should be noted that tourism is usually heavily linked with weather conditions. However, these factors explain less than 16% based on the overall R^2^. Detailed contributions of those factors can be found in the Appendix.

## 4. Discussion

### 4.1. Endemic species conservation: The role of feral dog management

On Navarino Island, feral dogs can choose between endemic species (mainly geese, such as the upland goose, and guanacos) or free-roaming as well as contained livestock (cows, horses, pigs) as prey [18], the latter with the risk of being disturbed or culled by people when attacking livestock. Since the number of invasive prey species exceeds that of endemic prey on Navarino Island, we expect the majority of their diet to consist of invasive species. This seems to be reflected in the low contribution of prey availability to predicted habitat suitability for feral dogs, as they have available food sources across the whole study area. Nevertheless, predation and disturbance of native species are highly likely. Given the significant population decline of guanacos over the past 20 years [24], with no natural predator in this ecosystem aside from dogs, predation is likely the main cause of this decline.

Nest predation by feral dogs likely exacerbates the documented nest predation by American minks on waterfowl [22]. We found that feral dog presence had a statistically significant impact on two-thirds of our focal species. In *Chloephaga*, feral dog occurrences and the likelihood of feral dog presence were significantly negatively associated with predicted *Chloephaga* occurrence. Feral dogs’ predicted preferences for habitats in forest regions, shrublands, as well as coastal areas and peat bogs, likely cause high niche overlap with waterfowl, especially geese. In contrast, flightless steamer ducks, which prefer breeding grounds in coastal areas and on islets, are less vulnerable to feral dog predation. This result is reflected in the model output, where dog occurrence had no significant effect on predicted flightless steamer duck occurrences.

In guanacos, the predicted likelihood of dog presence had a significant negative impact on predicted guanaco occurrence. This is particularly severe considering that guanacos and feral dogs had an estimated habitat overlap of over 38%, based on their presence records.

Feral dogs and village dogs are likely to interact. Wild canids are known to interbreed with domestic free-ranging dogs [44] [45]. We assume that feral dogs and free-ranging dogs, or village dogs, deliberately interact on Navarino Island. There have been observations of interbreeding between feral and village dogs on Navarino Island. However, our results suggest that feral dogs and village dogs show different patterns in habitat use, potentially due to their motivations and foraging strategies. While free-ranging village dogs follow tourists and beg for food [36], feral dogs seem to avoid interactions with humans. Considering that free-ranging village dogs are more likely to experience interactions with humans as positive, feral dogs plausibly have contrasting experiences where encounters with humans could lead to injuries or even death. Notwithstanding this, we cannot exclude the possibility of village dog samples being present among those samples we counted as feral dog samples.

Home ranges (HRs) of feral dogs are highly variable. Reported HRs range from 0.23 km^2^ in Puerto Natales, Chile [46] to 0.13 to 2.85 km^2^ in Puerto Rico [47]. Considering that we recorded 0.19 sightings per square kilometer in the 2514 km^2^ area of Navarino Island, feral dog density, as well as HRs, might be higher on Navarino Island. Most of our recordings (41.55%) were at the 275 km^2^ central area of the island, surrounded by mountains and lakes, serving as potential borders between habitats. We assume that the central area is the core area of at least one feral dog population. This is supported by the results of our MaxEnt prediction and the connectivity analysis, which showed that the central area, with the highest predicted habitat suitability, shows low connectivity to the east of the island but less restricted movement corridors to the south and west. This could indicate two or more separate dog populations in the east, center, and possibly west of the study area.

Factors such as sample size, environmental variable accuracy, and species’ ecological traits can affect model validity [30, 48]. Due to limited access to certain island regions, our sampling was uneven, with more reports from northern and central areas than from the southern region. This issue is common in presence-only models [49], where few samples come from regions with low animal presence. Additionally, double-counting cannot be avoided in occurrence-only sampling. Although we filtered our datasets to prevent double-counting feral dogs, we assume the actual number of upland geese and guanacos is drastically lower. The guanaco population is estimated to be a maximum of 20 individuals.

Our study was observational rather than experimental, so the patterns we observed could potentially be influenced by unrelated environmental factors. However, this is unlikely. Biophysical characteristics such as vegetation, soil types, topography, and rainfall, as well as protection from human disturbances, are consistent between areas with and without dogs in our study area. The presence of dogs can lead to changes in habitat use [50, 51]. In our study, upland geese did not exhibit different habitat use patterns between areas with and without dogs. However, guanacos seem to have shifted their habitat use to areas presumably less used by feral dogs, such as mountain ranges. Most guanaco sightings were on Isla Scott, an islet connected to Navarino Island by a small, regularly flooded land bridge. Sparsely vegetated areas and regions at risk of flooding might be less suitable for dogs and therefore serve as a refugium for small guanaco populations. This is supported by our MaxEnt predictions for feral dogs and guanacos, as well as the connectivity analyses of feral dog habitats. Although we did not observe predation events on guanacos nor record any evidence for dogs preying upon their carcasses, we cannot find any indication that guanacos are not disturbed by feral dogs at least, or more drastically, fall prey to them. Guanacos have not been sighted along the northern coast for many years [52], and their densities were as low as 0.14 individuals/km^2^ on the northeastern coast of Navarino Island during 2002–2005 [53]. Nowadays, they are presumably much lower, considering none or insufficient recorded reproduction rates. Thus, observing predation or harassment of guanacos by feral dogs is exceedingly rare. However, individual dog attacks on vulnerable species can substantially impact their persistence. Evidence of strong habitat overlap and recorded disturbances should raise awareness and prompt urgent conservation actions.

### 4.2 Feral dog classification: Impacts on conservation policies

Since 2012, the Servicio Agŕıcola y Ganadero (SAG) has implemented a standardised national protocol for addressing complaints related to wild carnivores attacking livestock. This protocol has facilitated the quantification of damages attributed to predatory dogs, effectively distinguishing their impacts from those caused by other wild carnivores.

In the Magallanes region, an evaluation of complaints over a 13-year period (2012–2023) documented a total of 123 complaints, averaging 10 per year, and involving 4,379 animals attacked, with a mean of 364 annually. Dogs were identified as responsible for 82.2% of the attacks, with 88.7% of these incidents involving sheep. Other affected livestock included poultry, cattle, and camelids. In contrast, wild carnivores such as foxes (7.0%) and pumas (4.3%) accounted for the remaining incidents.

Considering that feral dogs might also impact vulnerable and endangered birds of prey, management solutions are even more urgent. In Chile, discussions about lethal control of free-ranging dogs are complicated by legal ambiguities regarding whether a dog is classified as feral. Under Ley 19.473 (Ley de Caza), feral dogs are classified as part of the wildlife as they live freely and independently of humans. However, Decree 05/98 (the Regulation of the Law) does not include feral dogs in Article 6, which lists “harmful or injurious” species. Consequently, feral dogs are not subject to regulation under the hunting law.The Chilean Responsible Pet Ownership Law (Ley 21,020, Ministerio de Salud, Chile, 2017) defines several dog categories but does not recognise feral dogs. From a conservation perspective, this is a drastic oversight, as feral dogs threaten wildlife through predation and disturbance; for example, their presence on sandy beaches threatens shorebird populations [54]. They may also spread diseases to native mammals [55, 56] as well as transmit zoonoses to humans. Management strategies should focus on the impacts caused by dogs, not just on how they are categorised. Furthermore, dogs are the leading cause of livestock loss for small farmers, causing severe economic and animal welfare problems [57, 58]. Attacks by dogs on livestock (both large and small) in rural areas of Magallanes constitute a documented and objective issue, verified by the Agriculture and Livestock Service (SAG) within its routine technical activities. In addition to the measurable negative impacts on rural production and economy, these attacks significantly affect wildlife conservation and animal welfare.

The Chilean public shows a preference for non-lethal management strategies aligned with ethical considerations regarding animal treatment [59]. Non-lethal control methods, such as public education and mass neutering, may help limit long-term population growth but do not address the immediate threats posed by existing dog populations. Sterilised dogs still prey on wildlife, transmit diseases, and attack livestock. sterilisation does not reduce male dogs’ average roaming range [60], and in some cases, it increases their activity due to heightened food-seeking behavior [60], leading to increased predation.

Contreras-Abarca et al. [61] proposed the following criteria for classifying dogs (feral vs. owned) and as a management tool, based on criteria established by Boitani et al. [23] and Vanak and Gompper [51]:

Ownership status (owned/unowned): A dog is considered owned if it returns to one or more specific individuals who recognise the dog as their own.

1. Ranging behavior (restricted/free-ranging): A dog is considered restricted if its movement is limited to enclosed private property and it is supervised by a person when on public land.
2. Location (core area within human settlements/outside human settlements): Human settlements are defined as urban or rural areas with permanent human activity. Areas outside human settlements are regarded as wilderness.
3. To improve the safety of owned dogs and address feral dog impacts, we propose that all three criteria should be applied in combination when deciding whether a dog is feral.

To achieve immediate and effective results in improving current regulations in Chile, it is imperative to address the removal of free-roaming dogs in urban environments and feral dogs in rural areas. This would necessitate the implementation of lethal control methods as part of the management strategy.

### 4.3 Ecosystem transformation in the presence of invasive species

When comparing population size and species diversity with a vegetation-ecologically similar reserve in southern Chile, Torres del Paine National Park, clear differences emerge. No invasive predators had been recorded in Torres del Paine until recently, although the presence of American mink *(Neovison vison)* was considered likely. This assumption was confirmed in December 2024, when the first photographic evidence of mink was obtained in the Paine Grande area (Soto, 2024, personal communication).These differences support our findings, indicating a significant impact of feral dogs on endemic prey species and highlighting the importance of targeted management of invasive species, especially terrestrial predators, in the Cape Horn Biosphere Reserve.

Changes in NDVI over time likely result from a combination of factors, including climate change, increasing tourism, and shifts in species composition—especially the decline of endemic species alongside a rise in invasive ones. Among these, the proliferation of invasive beavers has a particularly notable impact on Patagonian ecosystems, with significant effects on native species like the Magellanic woodpecker. The increase of invasive beavers has led to significant shifts in Navarino Island’s vegetation through dam construction, which floods large areas and converts forested regions to wetlands, reducing the density of woody vegetation, disrupting nutrient cycling, and shifting the ecosystems toward a wetland-adapted flora [62, 63, 64]. Additionally, Anderson et al. [62] found that on Navarino Island in Chile, beaver activity drastically reduces riparian forest canopy cover, with effects reaching up to 30 metres from watercourses. This reduction severely impacts native tree species like *Nothofagus betuloides* and *Nothofagus pumilio*, while creating favourable conditions for exotic plants to invade the modified meadows. In this altered landscape, herbaceous species richness nearly doubled, but much of this increase was due to non-native species, leading to the replacement of native plant communities. Albeit guanacos as native ungulates might affect forest structure through browsing and trampling, which typically increases understory vegetation but reduces canopy cover, species richness, and diversity, resulting in a thinner leaf litter layer and potentially slowing forest succession [65], this impact cannot be found here. The decline in guanacos rather seems to lead to increased forestal vegetation, but a decrease in important habitat, which can displace native species and further degrade habitats that are critical for wildlife, such as ground-nesting birds.

The ecological dynamics on Navarino Island highlight alarming trends in the populations of invasive and native species, with beaver population numbers estimated at approximately 20,000 individuals in the early 2000s, based on observations, now occupying over 15% of Tierra del Fuego [66, 67], suggesting a similar trend on Navarino Island, where their numbers may have surged closely to carrying capacity. As Lizarralde Lizarralde et al. [64] and Skewes et al. [67] previously predicted, population growth is subject to ecological constraints, with a carrying capacity *K* of just over 100,000 for Tierra del Fuego and Isla Navarino combined. Once these populations reach a plateau, further exponential growth is unlikely. This underscores the importance of precise population monitoring and management to address the impact of invasive species. In the meantime, the guanaco population, once estimated at approximately 400 individuals, has dwindled to perhaps only a few dozen [42]. With no observed evidence of reproduction on Navarino Island, their risk of extinction appears increasingly imminent. With feral dogs contributing to a trophic cascade alongside the beavers and free-ranging farm animals, threatening endemic wildlife and the overall health of the ecosystem on Navarino Island.

In Torres del Paine, a sub-Antarctic ecosystem largely devoid of invasive species—particularly invasive predators—environmental conditions for the focal species are markedly more favourable than on Navarino Island. Habitat transformations in this region are more likely driven by over-arching climatic trends rather than direct anthropogenic pressures such as tourism or the introduction of non-native species. Guanacos, which show a preference for open steppe habitats, are already demonstrating population growth in Torres del Paine in contrast to Navarino Island, suggesting that the ecological shift toward steppe-dominated landscapes supports their long-term viability. In stark contrast, Navarino Island is experiencing an expansion of wetland areas, flooded areas and forest density alongside a decline in intermediate vegetation types. This, coupled with an increasing presence of predators, contributes to the degradation of habitat suitability for guanacos and other target species, rendering the island a considerably more challenging environment for their persistence.

An ecosystem dominated by invasive fauna and flora can lead to drastic habitat alterations that make these areas increasingly inhospitable for native wildlife. These invasive pressures may lead to homogenised ecosystems dominated by non-native species, further reducing biodiversity. Over time, this ecosystem transformation may result in a landscape that no longer supports the unique ecological niches required by endemic species, potentially leading to local extinctions and long-term ecosystem degradation.

## Limitations

The monitoring techniques had several limitations. Since the field expeditions on Navarino Island were limited to one area at a time and excluded paths that were impossible or too dangerous to access, the observed presences and absences have an unknown uncertainty. In Torres del Paine, guanacos and waterfowl could only be observed in the sight range, with individuals out of range potentially being unnoticed. Due to repeated observations, this error is minimised, but might still lead to a lower than actual estimated abundance.

## Author contributions

Hana Tebelmann: Conceptualisation, Methodology, Data curation, Formal analysis, Funding acquisition, Writing-Original draft preparation, Visualisation, Investigation, Validation, Writing-Reviewing and Editing. Alicia Sophie Pages: Data curation, Writing-Original draft preparation, Translation, Writing-Reviewing and Editing. Simon Käfer: Investigation, Formal analysis, Writing-Original draft preparation, Validation, Writing-Reviewing and Editing. Aintzane Cariñanos Ruiz: Data curation, species monitoring (Torres del Paine), Validation. Nicolas Soto Volkart: Data curation, formal analysis (Torres del Paine), Validation, Writing-Reviewing and Editing. Udo Gansloßer: Writing-Reviewing and Editing, Supervision.

## Limitations

The authors wish to note that a serious accident involving the expedition leader occurred during the second expedition. As a result, data collection could not continue for the accompanying team members. The expedition was not able to proceed continuously due to the accident and had to be paused during recovery. Immediately following the recovery, the expedition and data curation were resumed solely by the expedition leader.

## Acknowledgements

A significant part of the conceptualisation and data curation was inspired by the previous and current work of the dedicated researchers at UMAG and the sub-Antarctic Research Center in Puerto Williams, Chile. We would like to thank all the people involved, because without them, this project would not have been possible. Special thanks go to Michel Winter and Jantje Pasternak, who, unfortunately, were unable to continue the project after the accident. We are grateful for their support throughout the expedition and their willingness to collect data under such challenging conditions. Our deepest gratitude goes first and foremost to Pascal Guerin, as well as to the brave volunteer firefighters and Carabineros, for their rescue efforts during an expedition accident. Additionally, we would like to extend our heartfelt thanks to Pascal Guerin for his unwavering support and assistance, both during and after the expedition. We would like to thank Michael Thompson for his personal support during the two expeditions and for being an inspiration during bio-philosophical discussions. We would also like to thank Brenda Riquelme for her collegial and personal support. We would additionally like to thank Stella Gorrizizzo, Lena Albers, Olaf Bininda-Emonds, and Wilko Ahlrichs for their collegial assistance with the project; and Imke ter Fehr, Marcel Bartetzski, Tobias Herzog, Wiebke Meiners, Charlotte Kaluza, Malin Steindorf, and Marla Hiepler for their emotional encouragement and support.

We gratefully acknowledge Rab, Exped, Nordisk, Pulsar and Travellunch for supplying the expedition equipment used in our research. This support was limited to equipment only, with no financial contribution.

## Appendix

### Population growth estimation

To estimate population growth in guanacos and waterfowl, we used the following formula to estimate the growth rate for further statistical calculations.

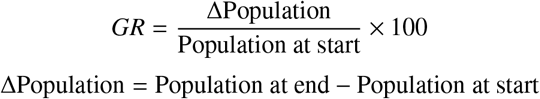

### Model selection

Models for predicted waterfowl occurrence were selected using lasso regression for variable reduction and parameter estimation. Only parameters with an estimated importance were included in the models. The model for predicted guanaco occurrence was selected based on the criteria included by the MaxEnt prediction and not further reduced, to account for a detailed analysis of variation in habitat selection.

### ARIMA output

Our ARIMA model consisted of autoregressive (AR) terms, differencing (I), and moving average (MA) components as well as seasonal AR, MA, and differencing terms. The fitted model was

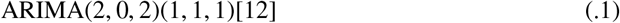

where we used 2 autoregressive and moving average terms, no differentiation term, since the data was stationary, as well as 1 seasonal AR, MA and differencing term to capture seasonal changes, which are likely present in sub-Antarctic regions.

#### Autoregressive terms (AR1, AR2)

These terms capture how past NDVI values influence future predictions. In our model:

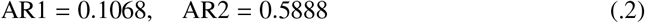

The moderate AR2 suggests shifts in NDVI over time. **Moving average terms (MA1, MA2)**: These account for past errors in predictions:

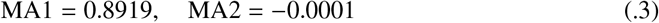

High MA1 means that past prediction errors significantly impact future NDVI estimates, so that a correction based on past prediction errors increases precision of future predictions. **Seasonal AR and MA (SAR1, SMA1)**: These capture annual cycles in NDVI:

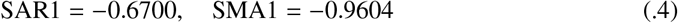

The high magnitude of SMA1 suggests strong seasonal patterns, meaning that NDVI fluctuations are strongly influenced by previous seasonal values, likely due to vegetation changes in winter and summer based on Antarctic climate. However, this does not account for gradual or subtle changes in habitat use and land cover, since the ARIMA builds predictions based on temporal patterns.

Residual diagnostics confirm the adequacy of the model. The Ljung-Box test:

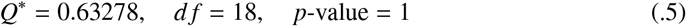

indicates no significant autocorrelation in residuals, suggesting a well-fitted ARIMA.

### Appendix. 1. NDVI modeling for Torres del Paine

## References

[1] S. H. Butchart, M. Walpole, B. Collen, A. Van Strien, J. P. Scharlemann, R. E. Almond, J. E. Baillie, B. Bomhard, C. Brown, J. Bruno, others, Global biodiversity: indicators of recent declines, Science 328 (2010) 1164–1168. Publisher: American Association for the Advancement of Science.

[2] H. Seebens, T. M. Blackburn, E. E. Dyer, P. Genovesi, P. E. Hulme, J. M. Jeschke, S. Pagad, P. Pyšek, M. Winter, M. Arianoutsou, others, No saturation in the accumulation of alien species worldwide, Nature communications 8 (2017) 14435. Publisher: Nature Publishing Group UK London.

[3] R. J. Hobbs, S. Arico, J. Aronson, J. S. Baron, P. Bridgewater, V. A. Cramer, P. R. Epstein, J. J. Ewel, C. A. Klink, A. E. Lugo, others, Novel ecosystems: theoretical and management aspects of the new ecological world order, Global ecology and biogeography 15 (2006) 1–7. Publisher: Wiley Online Library.

[4] M. E. Gompper, Introduction: outlining the ecological influences of a subsidized, domesticated predator, Free-ranging dogs and wildlife conservation (2014) 1–8. Publisher: Oxford University Press Oxford, UK.

[5] J. K. Young, K. A. Olson, R. P. Reading, S. Amgalanbaatar, J. Berger, Is wildlife going to the dogs? Impacts of feral and free-roaming dogs on wildlife populations, BioScience 61 (2011) 125–132. Publisher: American Institute of Biological Sciences Circulation, AIBS, 1313 Dolley ….

[6] J. Hughes, D. W. Macdonald, A review of the interactions between free-roaming domestic dogs and wildlife, Biological conservation 157 (2013) 341–351. Publisher: Elsevier.

[7] T. S. Doherty, C. R. Dickman, A. S. Glen, T. M. Newsome, D. G. Nimmo, E. G. Ritchie, A. T. Vanak, A. J. Wirsing, The global impacts of domestic dogs on threatened vertebrates, Biological conservation 210 (2017) 56–59. Publisher: Elsevier.

[8] W. M. Twardek, K. S. Peiman, A. J. Gallagher, S. J. Cooke, Fido, Fluffy, and wildlife conservation: The environmental consequences of domesticated animals, Environmental Reviews 25 (2017) 381–395. Publisher: NRC Research Press.

[9] E.A. Silva-Rodŕıguez, K. E. Sieving, Domestic dogs shape the landscape-scale distribution of a threatened forest ungulate, Biological Conservation 150 (2012) 103–110. Publisher: Elsevier.

[10] G. Zapata-ŕıos, L. C. Branch, Altered activity patterns and reduced abundance of native mammals in sites with feral dogs in the high Andes, Biological Conservation 193 (2016) 9–16. Publisher: Elsevier.

[11] A. T. Vanak, M. Thaker, M. E. Gompper, Experimental examination of behavioural interactions between free-ranging wild and domestic canids, Behavioral Ecology and Sociobiology 64 (2009) 279–287. Publisher: Springer.

[12] M. Clinchy, M. J. Sheriff, L. Y. Zanette, Predator-induced stress and the ecology of fear, Functional Ecology 27 (2013) 56–65. Publisher: Wiley Online Library.

[13] H. K. Epperly, M. Clinchy, L. Y. Zanette, R. A. McCleery, Fear of large carnivores is tied to ungulate habitat use: evidence from a bifactorial experiment, Scientific Reports 11 (2021) 12979. Publisher: Nature Publishing Group UK London.

[14] T. C. Wood, P. A. Moore, Fine-tuned responses to chemical landscapes: crayfish use predator odors to assess threats based on relative size ratios, Ecosphere 11 (2020) e03188. Publisher: Wiley Online Library.

[15] D. F. Sax, S. D. Gaines, Species invasions and extinction: the future of native biodiversity on islands, Proceedings of the National Academy of Sciences 105 (2008) 11490–11497. Publisher: National Acad Sciences.

[16] A. Sih, D. I. Bolnick, B. Luttbeg, J. L. Orrock, S. D. Peacor, L. M. Pintor, E. Preisser, J. S. Rehage, J. R. Vonesh, Predator–prey näıveté, antipredator behavior, and the ecology of predator invasions, Oikos 119 (2010) 610–621. Publisher: Wiley Online Library.

[17] F. Courchamp, J.-L. Chapuis, M. Pascal, Mammal invaders on islands: impact, control and control impact, Biological reviews 78 (2003) 347–383. Publisher: Cambridge University Press.

[18] E. Schüttler, L. Saavedra-Aracena, J. E. Jiménez, Domestic carnivore interactions with wildlife in the Cape Horn Biosphere Reserve, Chile: husbandry and perceptions of impact from a community perspective, PeerJ 6 (2018) e4124. Publisher: PeerJ Inc.

[19] J. Contardo, A. Grimm-Seyfarth, P. E. Cattan, E. Schüttler, Environmental factors regulate occupancy of freeranging dogs on a sub-Antarctic island, Chile, Biological Invasions 23 (2021) 677–691. Publisher: Springer.

[20] C. B. Anderson, C. R. Griffith, A. D. Rosemond, R. Rozzi, O. Dollenz, The effects of invasive North American beavers on riparian plant communities in Cape Horn, Chile: do exotic beavers engineer differently in sub-Antarctic ecosystems?, Biological Conservation 128 (2006) 467–474. Publisher: Elsevier.

[21] A. Maldonado-Márquez, T. Contador, J. Rendoll-Cárcamo, S. Moore, C. Pérez-Troncoso, D. Gomez-Uchida, C. Harrod, Southernmost distribution limit for endangered Peladillas (Aplochiton taeniatus) and non-native coho salmon (Oncorhynchus kisutch) coexisting within the Cape Horn biosphere reserve, Chile, Journal of Fish Biology 96 (2020) 1495–1500. Publisher: Wiley Online Library.

[22] E. Schüttler, R. Klenke, S. McGehee, R. Rozzi, K. Jax, Vulnerability of ground-nesting waterbirds to predation by invasive American mink in the Cape Horn Biosphere Reserve, Chile, Biological Conservation 142 (2009) 1450–1460. Publisher: Elsevier.

[23] L. Boitani, F. Francisci, P. Ciucci, G. Andreoli, J. Serpell, The ecology and behavior of feral dogs: a case study from central Italy, The domestic dog: its evolution, behaviour and interactions with people (2017) 342–368.

[24] B. González, ¿ Qué problemas de conservación tienen las poblaciones de guanaco en Chile, Ambiente forestal 5 (2010) 28–38.

[25] T. Václavík, R. K. Meentemeyer, Invasive species distribution modeling (iSDM): are absence data and dispersal constraints needed to predict actual distributions?, Ecological modelling 220 (2009) 3248–3258. Publisher: Elsevier.

[26] H. Costa, N. B. Ponte, E. B. Azevedo, A. Gil, Fuzzy set theory for predicting the potential distribution and cost-effective monitoring of invasive species, Ecological Modelling 316 (2015) 122–132. Publisher: Elsevier.

[27] L. Brotons, W. Thuiller, M. B. Araújo, A. H. Hirzel, Presence-absence versus presence-only modelling methods for predicting bird habitat suitability, Ecography 27 (2004) 437–448. Publisher: Wiley Online Library.

[28] D. I. MacKENZIE, What are the issues with presence-absence data for wildlife managers?, The Journal of Wildlife Management 69 (2005) 849–860. Publisher: Wiley Online Library.

[29] D. Fernández, M. Nakamura, Estimation of spatial sampling effort based on presence-only data and accessibility, Ecological Modelling 299 (2015) 147–155. Publisher: Elsevier.

[30] J. Elith, S. J. Phillips, T. Hastie, M. Dudík, Y. E. Chee, C. J. Yates, A statistical explanation of MaxEnt for ecologists, Diversity and distributions 17 (2011) 43–57. Publisher: Wiley Online Library.

[31] E. S. Bakker, H. Olff, M. Boekhoff, J. M. Gleichman, F. Berendse, Impact of herbivores on nitrogen cycling: contrasting effects of small and large species, Oecologia 138 (2004) 91–101. doi:10.1007/s00442-003-1402-5.

[32] F. Keesing, T. P. Young, Cascading consequences of the loss of large mammals in an african savanna, BioScience 64 (2014) 487–495. doi:10.1093/biosci/biu059.

[33] R. Rozzi, J. J. Armesto, J. R. Gutiérrez, F. Massardo, G. E. Likens, C. B. Anderson, A. Poole, K. P. Moses, E. Hargrove, A. O. Mansilla, others, Integrating ecology and environmental ethics: Earth stewardship in the southern end of the Americas, BioScience 62 (2012) 226–236. Publisher: American Institute of Biological Sciences Circulation, AIBS, 1313 Dolley ….

[34] E. Pisano, Fitogeografía de Fuego-Patagonia chilena (1977). Publisher: Instituto de la Patagonia Punta Arenas.

[35] F. M. Jaksic, J. A. Iriarte, J. E. Jiménez, The raptors of torres del paine national park, chile: biodiversity and conservation, Revista chilena de historia natural 75 (2002) 149–161. doi:10.4067/S0716-078×2002000200014.

[36] E. Schüttler, L. Saavedra-Aracena, J. E. Jiménez, Spatial and temporal plasticity in free-ranging dogs in sub-antarctic chile, Applied Animal Behaviour Science 250 (2022) 105610.

[37] D. L. Warren, S. N. Seifert, Ecological niche modeling in Maxent: the importance of model complexity and the performance of model selection criteria, Ecological applications 21 (2011) 335–342. Publisher: Wiley Online Library.

[38] A. Radosavljevic, R. P. Anderson, Making better Maxent models of species distributions: complexity, overfitting and evaluation, Journal of biogeography 41 (2014) 629–643. Publisher: Wiley Online Library.

[39] R. Anantharaman, K. Hall, V. B. Shah, A. Edelman, Circuitscape in Julia: High Performance Connectivity Modelling to Support Conservation Decisions, Proceedings of the JuliaCon Conferences 1 (2020) 58. doi:10.21105/jcon.00058, Publisher: The Open Journal.

[40] J. H. Friedman, T. Hastie, R. Tibshirani, Regularization Paths for Generalized Linear Models via Coordinate Descent, Journal of Statistical Software 33 (2010) 1–22. doi:10.18637/jss.v033.i01.

[41] M. E. Brooks, K. Kristensen, K. J. v. Benthem, A. Magnusson, C. W. Berg, A. Nielsen, H. J. Skaug, M. Mächler, B. M. Bolker, glmmTMB Balances Speed and Flexibility Among Packages for Zero-inflated Generalized Linear Mixed Modeling, The R Journal 9 (2017) 378–400. doi:10.32614/RJ-2017-066.

[42] B. A. González, et al., Maintenance of genetic diversity in an introduced island population of guanacos after seven decades and two severe demographic bottlenecks: Implications for camelid conservation, PLOS ONE 9 (2014) e91714. doi:10.1371/journal.pone.0091714.

[43] NASA POWER Project, NASA POWER Data Access Viewer, 2024. https://power.larc.nasa.gov.

[44] A. Caragiulo, S. J. Gaughran, N. Duncan, C. Nagy, M. Weckel, B. M. vonHoldt, Coyotes in new york city carry variable genomic dog ancestry and influence their interactions with humans, Genes 13 (2022) 1661. doi:10.3390/genes13091661.

[45] N. Kopaliani, M. Shakarashvili, Z. Gurielidze, T. Qurkhuli, D. Tarkhnishvili, Gene flow between wolf and shepherd dog populations in georgia (caucasus), Journal of Heredity 105 (2014) 345–353.

[46] G. E. Pérez, A. Conte, E. J. Garde, S. Messori, R. Vanderstichel, J. Serpell, Movement and home range of owned free-roaming male dogs in Puerto Natales, Chile, Applied Animal Behaviour Science 205 (2018) 74–82. doi:10.1016/j.applanim.2018.05.022.

[47] C.C. Sauvé, A. R. Berentsen, S. F. Llanos, A. T. Gilbert, P. A. Leighton, Home range overlap between small Indian mongooses and free roaming domestic dogs in Puerto Rico: implications for rabies management, Scientific Reports 13 (2023) 22944. URL: 10.1038/s41598-023-50261-7. doi:10.1038/s41598-023-50261-7.

[48] P. A. Hernandez, C. H. Graham, L. L. Master, D. L. Albert, The effect of sample size and species characteristics on performance of different species distribution modeling methods, Ecography 29 (2006) 773–785. Publisher: Wiley Online Library.

[49] C. B. Yackulic, R. Chandler, E. F. Zipkin, J. A. Royle, J. D. Nichols, E. H. Campbell Grant, S. Veran, Presence-only modelling using MAXENT: when can we trust the inferences?, Methods in Ecology and Evolution 4 (2013) 236–243. Publisher: Wiley Online Library.

[50] E.A. Silva-RodŔiguez, G.R. Ortega-Soíis, J. E. Jiménez, Conservation and ecological implications of the use of space by chilla foxes and free-ranging dogs in a human-dominated landscape in southern Chile, Austral Ecology 35 (2010) 765–777. Publisher: Wiley Online Library.

[51] A. T. Vanak, M. E. Gompper, Dogs Canis familiaris as carnivores: their role and function in intraguild competition., Mammal review 39 (2009).

[52] B. González, B. Zapata, J.C. Maŕın, Situación de conservación y percepción local sobre la población de guanacos más austral del mundo, Isla Navarino, XII Región de Chile, Report, Pontificia Universidad Católica de Chile, 2002.

[53] B. A. González, Informe sobre la situación de conservación de la población de guanacos (Lama guanicoe) en la costa Noroeste de isla Navarino, XII Región de Chile, Report, Fundación Biodiversitas, Chile, 2005.

[54] E. I. Cortés, J. G. Navedo, E.A. Silva-Rodŕıguez, Widespread presence of domestic dogs on sandy beaches of southern Chile, Animals 11 (2021) 161. Publisher: MDPI.

[55] E. Hidalgo-Hermoso, J. Cabello, C. Vega, H. Kroeger-Gómez, D. Moreira-Arce, C. Napolitano, C. Navarro, I. Sacristán, A. Cevidanes, S. Di Cataldo, others, An eight-year survey for canine distemper virus indicates lack of exposure in the endangered Darwin’s fox (Lycalopex fulvipes), The Journal of Wildlife Diseases 56 (2020) 482– 485. Publisher: Wildlife Disease Association.

[56] S. Llanos-Soto, D. González-Acuña, Conocimiento acerca de los patógenos virales y bacterianos presentes en mamíferos silvestres en Chile: una revisión sistemática, Revista chilena de infectología 36 (2019) 43–67. Publisher: SciELO Chile.

[57] D. Montecino-Latorre, W. San Martín, Evidence supporting that human-subsidized free-ranging dogs are the main cause of animal losses in small-scale farms in Chile, Ambio 48 (2019) 240–250. Publisher: Springer.

[58] J. Butler, J. du Toit, J. Bingham, Free-ranging domestic dogs (Canis familiaris) as predators and prey in rural Zimbabwe: threats of competition and disease to large wild carnivores, Biological Conservation 115 (2004) 369– 378. doi:10.1016/S0006-3207(03)00152-6.

[59] F. J. Villatoro, L. Naughton-Treves, M. A. Sepúlveda, P. Stowhas, F. O. Mardones, E.A. Silva-Rodŕıguez, When free-ranging dogs threaten wildlife: Public attitudes toward management strategies in southern Chile, Journal of Environmental Management 229 (2019) 67–75. doi:10.1016/j.jenvman.2018.06.035.

[60] E. Garde, G. E. Pérez, R. Vanderstichel, P. F. D. Villa, J. A. Serpell, Effects of surgical and chemical sterilization on the behavior of free-roaming male dogs in Puerto Natales, Chile, Preventive Veterinary Medicine 123 (2016) 106–120. doi:10.1016/j.prevetmed.2015.11.011.

[61] R. Contreras-Abarca, S. J. Crespin, D. Moreira-Arce, J. A. Simonetti, Redefining feral dogs in biodiversity conservation, Biological Conservation 265 (2022) 109434. doi:10.1016/j.biocon.2021.109434.

[62] C. B. Anderson, C. R. Griffith, A. D. Rosemond, R. Rozzi, O. Dollenz, The effects of invasive north american beavers on riparian plant communities in cape horn, chile: Do exotic beavers engineer differently in sub-antarctic ecosystems?, Biological Conservation 128 (2006) 467–474. doi:10.1016/j.biocon.2005.10.011.

[63] C. B. Anderson, A. D. Rosemond, Ecosystem engineering by invasive exotic beavers reduces in-stream diversity and enhances ecosystem function in cape horn, chile, Oecologia 154 (2007) 141–153. doi:10.1007/s00442-007-0757-4.

[64] M. S. Lizarralde, G. Deferrari, S. E. Alvarez, A long-term study on the impact of north american beaver on native trees in tierra del fuego, Biological Conservation 119 (2004) 565–573. doi:10.1016/j.biocon.2003.12.015.

[65] J. I. Ramirez, P. A. Jansen, J. den Ouden, L. Goudzwaard, L. Poorter, Long-term effects of wild ungulates on the structure, composition and succession of temperate forests, Forest Ecology and Management 432 (2019) 478–488. doi:10.1016/j.foreco.2018.09.049.

[66] C. B. Anderson, G. MartínezPastur, M. V. Lencinas, P. K. Wallem, M. C. Moorman, A. D. Rosemond, Do introduced north american beavers Castor canadensis engineer differently in southern south america? an overview with implications for restoration, Journal of Applied Ecology 45 (2008) 1230–1238. doi:10.1111/j.1365-2907.2008.00136.x.

[67] O. Skewes, C. Salgado, J. Yáñez, J. Bertram, Abundance and distribution of american beaver, Castor canadensis (kuhl 1820), in tierra del fuego and navarino islands, chile, European Journal of Wildlife Research 52 (2006) 292–296. doi:10.1007/s10344-006-0038-2.

